# HP1 proteins compact DNA into mechanically and positionally stable phase separated domains

**DOI:** 10.1101/2020.10.30.362772

**Authors:** Madeline M. Keenen, David Brown, Lucy D. Brennan, Roman Renger, Harrison Khoo, Christopher R. Carlson, Bo Huang, Stephan W. Grill, Geeta J. Narlikar, Sy Redding

## Abstract

In mammals HP1-mediated heterochromatin forms positionally and mechanically stable genomic domains even though the component HP1 paralogs, HP1*α*, HP1*β*, and HP1*γ*, display rapid on-off dynamics. Here we investigate whether phase-separation by HP1 proteins can explain these biological observations. Using bulk and single-molecule methods, we show that, within phase-separated HP1*α*-DNA condensates, HP1*α* acts as a dynamic liquid, while compacted DNA molecules are constrained in local territories. These condensates are resistant to large forces yet can be readily dissolved by HP1*β*. Finally, we find that differences in each HP1 paralog’s DNA compaction and phase-separation properties arise from their respective disordered regions. Our findings suggest a generalizable model for genome organization in which a pool of weakly bound proteins collectively capitalize on the polymer properties of DNA to produce self-organizing domains that are simultaneously resistant to large forces at the mesoscale and susceptible to competition at the molecular scale.

## I. INTRODUCTION

Compartmentalization of the eukaryotic genome into active and repressed states is critical for the development and maintenance of cell identity[1, 2]. Two broad classes of genome compartments are heterochromatin, which contains densely packed DNA regions that are transcriptionally repressed, and euchromatin, which contains physically expanded DNA regions that are transcriptionally active[3–5]. A highly conserved type of heterochromatin involves the interaction of proteins from the heterochromatin Protein 1 (HP1) family with chromatin that is methylated on histone H3 at lysine 9[6–9]. In addition to repressing transcription, this type of heterochromatin also plays critical roles in chromosome segregation and in conferring mechanical rigidity to the nucleus[3, 10].From investigations of chromatin in cells, it is not immediately obvious how to connect the biophysical properties of HP1 proteins to the diverse roles of HP1-mediated heterochromatin. Heterochromatin domains are typically found to be statically positioned within the nucleus for several hours, held separate from euchromatin[11, 12].Yet these domains can also be rapidly disassembled in response to environmental and developmental cues[13–15]. The finding that HP1 molecules in these domains exchange within seconds provides some insight into how these domains can be dissolved, because competing molecules would be able to rapidly displace HP1 proteins from DNA[16, 17]. However, such models raise the fundamental question of how HP1 molecules, which are dynamic on the order of seconds, enable chromatin states that are stable on the order of hours, and further how these states can resist the forces exerted on chromatin in the cell. The mammalian genome contains three HP1 paralogs: HP1*α*, HP1*β* and HP1*γ*. While the three paralogs show a high degree of homology, they are associated with distinct biological roles[18, 19]. For example, HP1*α* is mostly associated with gene repression and chromosome segregation[3, 18, 19], HP1*β* plays both gene activating and gene repressive roles[3, 18, 19], and HP1*γ* is more often associated with promoting transcription[3, 18, 19]. These observations raise the question of how small differences at the amino acid level give rise to distinct biophysical properties that direct the different functions of the HP1 paralogs. Some of the questions raised above have been investigated *in vitro*. For example, it has been shown that HP1 proteins are sufficient to bind to DNA and chromatin and to provoke their robust condensation[20–26]. These experiments have led to a model where HP1 molecules, by means of multiple contacts, condense and staple chromatin structures in place.

Furthermore, and consistent with cellular measurements, HP1 molecules also exhibit weak affinity for chromatin *in vitro*[22, 27]. Recent findings of phase-separation behavior by HP1 proteins provide an added perspective to the questions above [24, 28–30]. Specifically, the human HP1 protein, HP1*α* was shown to undergo liquid-liquid phase separation (LLPS) *in vitro* when phosphorylated and in combination with DNA[24]. Parallel studies showed that the Drosophila HP1 protein, HP1a, also forms phase-separated condensates *in vivo*[29]. In contrast, HP1*β* cannot undergo LLPS *in vitro* upon phosphorylation or in combination with DNA, but can be recruited to liquid phases of modified chromatin[24, 30]. The biophysical interactions that give rise to *in vitro* LLPS are consistent with the *in vivo* observations of low affinity binding and chromatin condensation by HP1*α*. The weak interactions underlying HP1-mediated LLPS also provide an attractive rationale for the rapid invasion and disassembly of heterochromatin. However, such an LLPS-based model does not easily explain the mechanical and temporal stability of chromatin domains. A recent study has implied that HP1-mediated heterochromatin in cells does not exhibit liquid-like phase-separated behavior[31]. This conclusion was drawn from the material properties of a subset of LLPS systems *in vitro*, such as impermeable boundaries and concentration buffering. However, these properties do not translate simply from *in vitro* to *in vivo* settings as condensates in cells span a diversity of protein environments and solvation conditions that will vary the nature of their boundaries and partitioning of nuclear material. Such narrow definitions are not generally applicable and fail to capture the nature of several types of condensates[32, 33]. Specifically, for condensates that involve DNA, there are additional constraints that arise from the properties of long polymers that do not scale in a straightforward way from smaller systems. These important considerations underscore the need to move beyond simple definitions and better understand the different and sophisticated ways in which condensates play biological roles. Here, using a combination of ensemble and single-molecule methods, we uncover the molecular basis of intramolecular DNA compaction by HP1*α* and the molecular determinants that give rise to HP1*α*-induced phase separation. In doing so, we investigate the role of DNA in condensates, both as a binding partner for HP1*α* and as a long polymer with unique organizational constraints. We show that condensates of HP1*α* and DNA are maintained on the order of hours by HP1*α* binding that is dynamic on the order of seconds. We find that the central disordered region of HP1*α* is sufficient to enable LLPS with DNA, and that the additional disordered regions regulate the activity of this central region. These results are then leveraged to uncover intrinsic biophysical differences across the three human HP1 paralogs. Finally, we show that the HP1*α*-DNA condensates are resistant to mechanical disruption by large forces and yet can be readily dissolved by HP1*β*. Overall our results uncover specific biophysical properties of each HP1 paralog in the context of DNA that have general implications for interpreting and understanding the behaviors and functions of HP1 in the context of chromatin.

## II. RESULTS

From previous work, we’ve found that HP1*α* shows the most robust phase-separation and DNA compaction abilities of all of the HP1 paralogs[24]. We therefore first used HP1*α* and DNA as a model system to dissect the steps involved in DNA compaction and phase-separation and to study the material properties of the resultant phases. We then carried out structure-function analysis on HP1*α* to understand how different regions of HP1*α* contribute to phase-separation. The results from these studies provided a well-defined biophysical framework within which to (i) compare the activities of HP1*β* and HP1*γ*, and (ii) understand how HP1*β* and HP1*γ* impact the phase-separation activities of HP1*α*. Finally, throughout we compare our observations of HP1-DNA condensates with prevailing views of the expected behavior of condensates.

### A. HP1*α* binds DNA globally but compacts DNA locally

We have previously shown that HP1*α* rapidly compacts long stretches of DNA[24]. To understand the mechanism of DNA compaction, we have leveraged a single molecule DNA curtain approach (Figure 1A)[34]. In this assay, ~50kbp molecules of DNA from bacteriophage *λ* are fixed to the surface of a microfluidic flowcell via a supported lipid bilayer. Visualization of DNA is achieved by labeling with the intercalating dye YOYO-1 (Figure 1B-D,F). HP1*α* is then pulsed into the flowcell, driving rapid DNA compaction (Figure 1B,D-F, figure supplement 1.1A-C). Previously, we showed that HP1*α*induced DNA compaction is an electrostatically driven process that proceeds by first concentrating DNA at the free end, and then rapidly and sequentially incorporating upstream DNA into a single condensate (Figure 1B)[24]. We validated that compaction occurs at the free end by labeling the untethered end of the DNA with a fluorescent dCas9 (Figure 1C-E).

**Figure 1.**
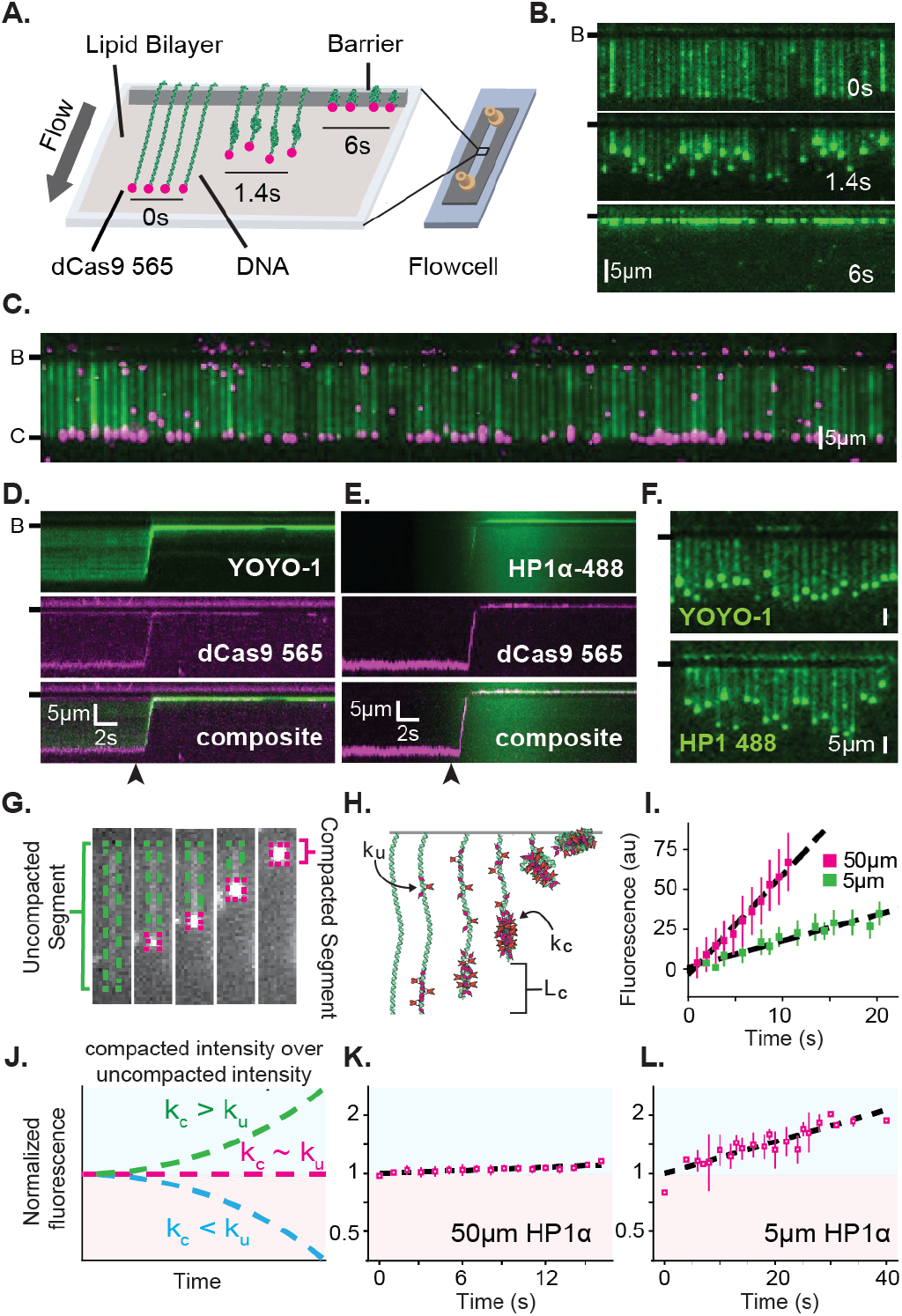
Characterization of DNA compaction by HP1*α*. **A.** Cartoon of the DNA curtains assay showing compaction of DNA. **B.** Timestamped images of DNA labeled with YOYO-1 undergoing compaction by 50*µ*M HP1*α* (unlabeled) shown before, during, and after compaction. (B-) or (-) specifies location of the barrier. **C.** DNA curtain end-labeled with fluorescent dCas9 (C-). The dCas9 is targeted to a site 750bp from the untethered end of the DNA. D. and E. Kymograms of DNA compaction by 50*µ*M HP1*α*. **D.** DNA labeled with YOYO-1 (top), dCas9-565 (middle), and composite image (bottom). **E.** HP1*α*-488 (top), DNA labeled with dCas9-565 (middle), and composite image (bottom). Arrowheads represent estimated time of protein injection. **F.** Still images during DNA compaction of either DNA labeled with YOYO-1 (top) or HP1*α*-488 (bottom). **G.** A DNA molecule undergoing compaction by HP1*α* specifying the uncompacted segment (green) and compacted segment (magenta). **H.** Cartoon of HP1*α* compacting DNA over time. *Lc* is the length of compacted DNA, *ku* is the rate of fluorescence increase for the uncompacted DNA segment, and *kc* is the rate fluorescence increase for the compacted DNA segment. See Materials and Methods for more information. **I.** Fluorescence increase of HP1*α*-488 on uncompacted DNA. **J.** Cartoon showing potential results from normalizing the fluorescence of the compacted segment by the uncompacted segment. **K.** and **L.** Measured normalized compacted HP1*α* fluorescence relative to uncompacted HP1*α*.

To further understand how HP1*α* compacts DNA, we directly visualized fluorescently labeled HP1*α* binding to DNA during compaction. Surprisingly, we found that HP1*α* binds uniformly along DNA, incorporating into both the compacted and uncompacted regions (Figure 1F-L). We observed a linear increase in fluorescence due to HP1*α* binding on uncompacted DNA (Figure 1I). And by comparison, we found that HP1*α* incorporates into compacted DNA at the same rate as on uncompacted DNA at 50*µ*M HP1*α*, and moderately faster into the compacted DNA at 5*µ*M HP1*α* (Figure 1J-L). We conclude that compacted DNA states are not inaccessibly compacted, but rather continue to support ingress and egress of HP1*α* from solution.

We considered two possibilities to explain how global binding would manifest in local compaction. In the first possibility HP1*α* binding is coupled to bending of the binding site. In such a case, the cumulative effect of multiple HP1*α* binding events would appear as a scrunching of the DNA fiber that would be evident in the fluorescence HP1*α* or DNA signal. However, we observe no appreciable increase in the YOYO-1 signal on noncompacted DNA during compaction (figure supplement 1.2A). In addition, the linear increase in fluorescence, due to HP1*α* binding, on the uncompacted segment of the DNA (Figure 1I) is consistent with HP1*α* binding in the absence of appreciable DNA bending of the binding site. Whereas a quadratic increase in HP1*α* fluorescence would be expected if the fluorescent signal was the product of HP1*α* association and increased local DNA density as a result of bending.

In the second possibility, HP1*α* molecules could trap naturally occurring DNA fluctuations by binding to multiple distal DNA sites simultaneously, or through the interactions of two or more HP1*α* molecules pre-bound to distal DNA sites. Indeed, the rapid and constant speed of DNA compaction against buffer flow (47kbp/s at ¡1pN for 50*µ*M HP1*α*) suggests that HP1*α* capitalizes upon DNA fluctuations that bring linearly distal segments of DNA together[35, 36]. Such a model then explains why the initiation of compaction is localized to the untethered end of the DNA: the lower tension at the untethered end allows for a larger number of DNA conformations that bring distal regions of the DNA into close proximity. HP1*α* is then able to trap these conformations leading to increased inclusion into the growing condensate either through HP1*α*-DNA or potentially through HP1*α*-HP1*α* interactions. The uniform binding of DNA by HP1*α* may additionally result in DNA that is easier to compact by altering the effective persistence length of the coated polymer.

From the results above, we identify three regulatable steps of HP1*α*-DNA condensation: local assembly of HP1*α* along DNA prior to DNA condensation, initiation of DNA compaction through capturing of lateral DNA fluctuations, and progression of DNA compaction through inclusion of uncompacted DNA into the growing condensate via HP1*α*-DNA and HP1*α*-HP1*α* interactions. As described in the discussion, nucleosomes and other nuclear factors will modulate each of these steps to further regulate DNA compaction.

### B. Condensate formation is more sensitive to the concentration of HP1*α* than of DNA

HP1*α* behaviors that result in DNA compaction at the single molecule level will also produce meaningful effects at the meso-scale. To further uncover the molecular details of how HP1*α* organizes DNA, we generated a phase diagram of HP1*α*-DNA condensation using short (147 bp) double stranded DNA oligomers (Figure 2A). The length of the DNA (near the persistence length for BDNA) was constrained to study the role of HP1*α*-DNA and potential HP1*α*-HP1*α* interactions while minimizing extensive polymer behaviors of DNA. At the conditions these experiments were performed (70 mM KCl, 20mM HEPES pH 7.5, 1mM DTT), HP1*α* remains soluble even at exceedingly high concentrations (400 *µ*M) (Figure 2A, bottom right panel). However, in the presence of DNA, HP1*α* readily condenses into concentrated liquid phase-separated material (Figure 2A) indicating the formation of a network of weak interactions interconnecting HP1*α* and DNA molecules. Such interactions are consistent with HP1*α*’s ability to capture and stabilize distal segments of DNA leading to DNA compaction as discussed in the previous section.

**Figure 2.**
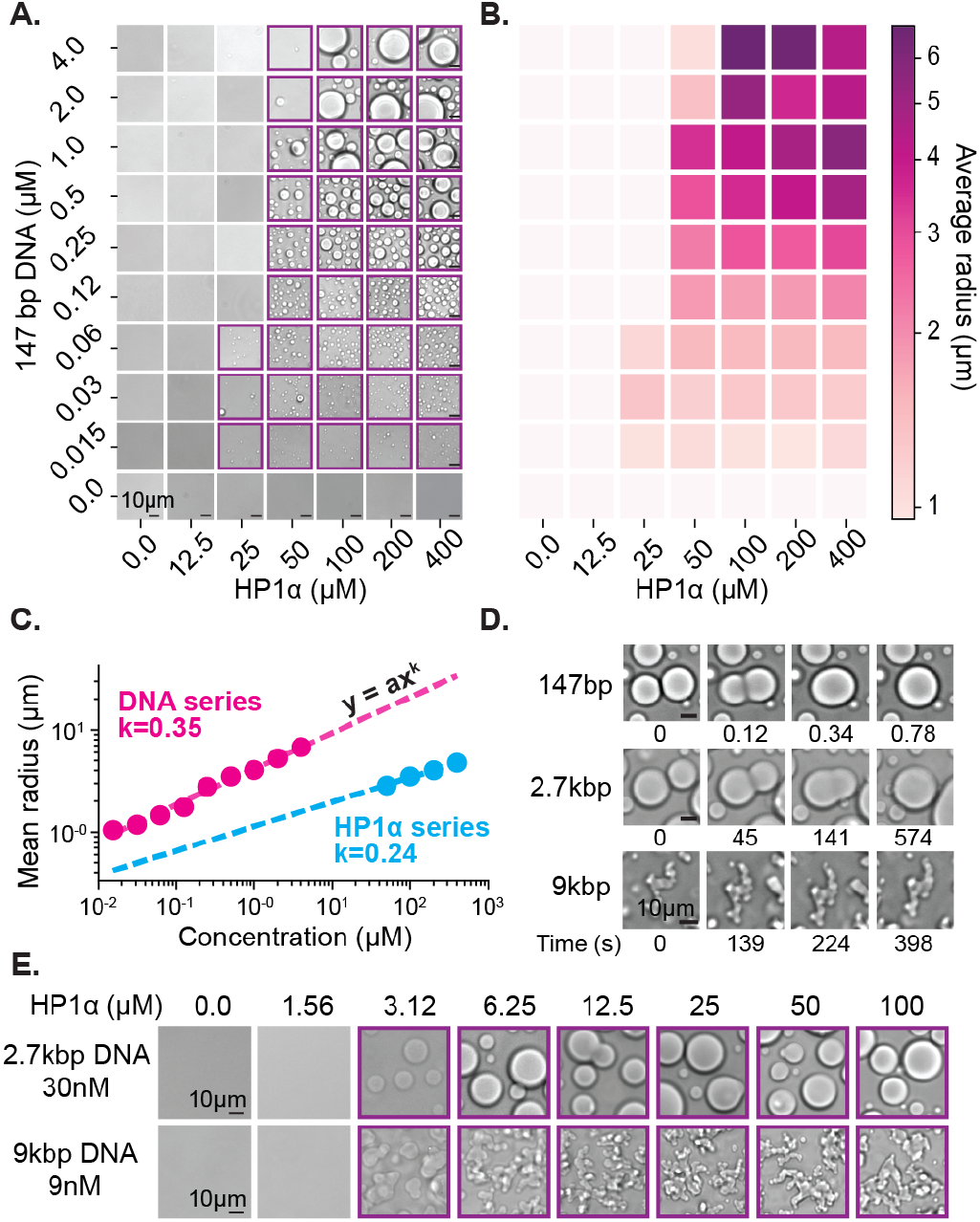
Characterization of HP1*α*-DNA condensate formation. **A.** Brightfield images of mixtures of HP1*α* and 147bp DNA. **B.** Heat map of the average radius of condensates for each condition in (A). **C.** Average condensate radius plotted against HP1*α* (cyan) or 147bp DNA (magenta) concentration and fit to a power law. **D.** Time stamped brightfield images of 100*µ*M HP1*α* and 147bp, 2.7kbp, or 9kbp DNA depicting fusion and coalescence behavior. **E.** Brightfield images of HP1*α* with either 30nM 2.7kb DNA (top) or 9nM 9kbp DNA (bottom). Throughout, purple boxes indicate presence of condensates.

One way to quantify the phase-separation capability of a molecule is through measurement of its critical concentration. Empirically, the critical concentration is defined as the concentration of the molecule above which the system separates into two phases. Theoretically, this transition occurs at the concentration at which the collective weak interactions of the system pay the entropic cost of de-mixing. In a two-component system, such as HP1*α* and DNA, each component may contribute differentially to condensation, and measuring the critical concentration of each component can provide insights into how the two components interact to form condensates.

First, we estimated the critical concentration of HP1*α* necessary to induce phase separation to be 50uM in the presence of 147bp DNA at concentrations ranging from 0.125uM to 4uM (Figure 2A). However, above this critical concentration of HP1*α*, we were unable to measure a corresponding critical concentration for DNA. Rather, lowering the DNA concentration resulted in a continuous reduction in the average size of observed HP1*α*-DNA condensates instead of a sharp disappearance (Figure 2A-C, figure supplement 2.1A). Thus, we conclude the critical concentration of HP1*α* is largely invariant of DNA concentration—even at sub-stochiometric ratios of DNA to HP1*α* (1:6000, figure supplement 2.2A).

The apparent equilibrium constant for HP1*α* interactions with 60-200 bp DNA ranges from 0.3-10 *µ*M[37](c.f. figure supplement 2.2D) which means, for most of the conditions tested here where we observe macroscopic droplets, we expect that nearly all DNA molecules are fully bound by HP1*α*. Once a collection of HP1*α* molecules coat a single DNA, that DNA molecule and its associated HP1*α* can, on average, act as a single highly valent molecule, or proto-condensate, that acts as a liquid building block and aggregates with other HP1*α*DNA proto-condensates as they encounter one another in solution[38]. It is helpful to recall that DNA regions already bound by HP1*α* were readily incorporated into condensates in our curtain assay, and the same biophysical considerations above also apply here. Specifically, we expect that condensate formation and growth are dependent on the concentration of HP1*α* and are the result of either higher order HP1*α* oligomerization or molecular rearrangements along DNA oligomers interacting in trans.

The ensuing aggregation process—proto-condensates clustering into large macroscopic condensates—should result in condensates sizes distributed according to a power law; where the power is set by molecular rates of diffusion and absorption[39–41]. Specifically, this result comes about because increasing the HP1*α* or DNA concentration increases the rate of formation and total number of proto-condensates, which increases their encounter frequency in solution accelerating the process of diffusion-driven aggregation. To test this hypothesis, we measured the average radius of condensates as a function of DNA and HP1*α* concentration (Figure 2B-C, figure supplement 2.1A). We find the average droplet size versus concentration of both DNA and HP1*α* is in fact well described by a power law (Figure 2C), further connecting the formation of macroscopic liquid droplets to the microscopic processes of aggregation and DNA compaction. While our data are consistent with HP1*α*-DNA binding promoting higher order HP1*α* oligomerization, at the same time, prior work suggests that the interface involved in HP1*α*-HP1*α* interactions following phosphorylation overlaps with the interface involved in HP1*α*-DNA interactions. If HP1*α* oligomerization is a key factor driving condensation, we then predict that as DNA concentration is increased, eventually HP1*α*-DNA interactions will outcompete HP1*α*-HP1*α* interactions, resulting in a loss of condensation. However, an alternative, compatible explanation suggests that as DNA concentration is increased, each DNA molecule is no longer bound by a sufficient amount of HP1*α* to create a productive proto-condensate or stabilize macroscopic condensates. Consistent with both of these expectations, at concentrations approaching equimolar ratios of HP1*α* to DNA binding sites (assuming 60bp per HP1*α* dimer binding site (materials and methods)—At 50*µ*M HP1*α* and 2-4*µ*M 147bp DNA) droplet formation is abrogated (Figure 2A-B, figure supplement 2.2A).

Overall, the behavior of HP1*α* and DNA in this condensation assay is consistent with the compaction process we measure in our single molecule assay, and ultimately our results demonstrate that DNA and HP1*α* play qualitatively different roles in the formation of the HP1*α*-DNA condensates. In both assays, at suitable HP1*α* concentrations, HP1*α* condenses locally around a single DNA molecule. In the curtains assay, DNA is then compacted through lateral HP1*α*-DNA and possible HP1*α*-HP1*α* interactions in cis, whereas in the droplet assay, HP1*α* and DNA collectively condense into protoand macroscopic condensates in trans. Additionally, both assays suggest that HP1*α* engaged with a single DNA molecule samples the same biophysical states as HP1*α* molecules contained within compacted structures and large macroscopic phases. However, a key difference between these two assays is the length of DNA. We observe robust DNA condensation on curtains at concentrations lower than the critical concentration for HP1*α*-DNA LLPS measured here on short DNA oligomers (figure supplement 1.1B-C, Figure 2A), indicating changes in DNA length will affect the formation of condensates. Moreover, we expect that as DNA length is increased, the conformational constraints and increased binding site availability of longer polymers will also have profound effects on the formation and material properties of HP1*α*-DNA condensates.

### C. The length of the DNA affects critical concentration and viscosity

The above studies were designed to minimize the contributions of DNA polymer length to allow us to investigate how multivalent interactions between HP1*α* and DNA promote the formation of condensates. At the scale of individual HP1*α* molecules, these multivalent interactions have many similarities to the types of multivalent interactions described in liquid-liquid phase-separating protein-protein and protein-RNA systems[42, 43]. However, at genomic scales, two features of HP1*α*-DNA condensates are expected to diverge from other commonly studied phase-separating systems. First, the size disparity between DNA in the nucleus and HP1*α* is several orders of magnitude. Therefore, neither the valency nor concentration of DNA is expected be limiting for HP1*α* condensation in the nucleus. In contrast, conditions are possible in the cell where the valency and concentration of scaffolding RNA molecules or client proteins are in short supply. Second, the length of genomic DNA will have profound bulk-level effects on condensate viscosity and morphology that will be distinct from other phase separating biological mixtures. Consequently, current definitions need to be modified when discussing phases formed in the context of HP1 proteins to explicitly include the polymer behavior of DNA. Towards this goal, we next investigated the effects of increasing DNA length on HP1*α*-DNA condensates. We expected to observe two results: lower critical concentrations of HP1*α* necessary to induce condensation due to increases in DNA valency and increases in bulk viscosity resulting in subsequent changes to the shapes of condensates.

Upon increasing the size of linear DNA co-incubated with HP1*α* from 147bp to 2.7kbp, we observed an order of magnitude decrease in the critical HP1*α* concentration required to induce LLPS (50*µ*M to 3*µ*M) (Figure 2A,E). This result is consistent with the roughly one order of magnitude increase in estimated HP1*α* binding sites from 2 to 45 per DNA molecule[44]. However, further increasing the DNA length to 9kbp did not lead to an additional decrease in the critical HP1*α* concentration (Figure 2F). This outcome is interesting because the apparent lower limit we measure for the critical HP1*α* concentration is coincident with estimates of the HP1*α*HP1*α* dimerization constant[24]. This may mean that HP1*α* dimerization either increases DNA binding affinity, or that dimerization plays a specific role in the formation of condensates. In our single molecule assay, we observed HP1*α*-induced DNA compaction at concentrations as low as 500nM (figure supplement 1.1B-C). However, the rate of DNA compaction exhibited by 500nM HP1*α* was roughly 30 times slower than the compaction rate at 5uM HP1*α* where we might have predicted only a 10 times slower rate of compaction based on an expected change in the pseudo-first order association rate constant (figure supplement 1.1C). This suggests that HP1*α* dimerization modestly increases HP1*α*’s on-rate for DNA binding. In addition, the sharp loss of condensates at concentrations where DNA binding and slower DNA compaction still occurs, indicates that dimerization is kinetically upstream of condensate formation and/or affects HP1*α*-DNA binding parameters, which are not critical during compaction.

In addition to changes in critical concentration, we also inear, which has been confirmed experimentally[45, 46]. Thus, the increase in size of linear DNA from 147bp to 2.7kbp should approximately correspond to an order of magnitude change in viscosity. However, while coalescence was complete within one second for condensates formed with 147bp DNA, condensates formed withobserve a marked reduction in the rate of coalescence of HP1*α*-DNA condensates formed from longer DNA lengths (Figure 2D). HP1*α*-DNA condensates formed with 147bp DNA rapidly coalesce into spherical structures immediately following fusion (Figure 2D). However, increasing the DNA length to 2.7kbp substantially (¿100times) lengthens the time required for coalescence (Figure 2D). Such slower coalescence could be reflective of decreasing surface tension and/or increasing viscosity. It is unlikely that DNA-DNA binding modes contribute to the condensate surface tension. Therefore, we assume that surface tension arises through HP1*α*-DNA and potentially HP1*α*-HP1*α* interactions, which should both be unchanged in character upon increasing DNA length. Instead we expect that the increased intrinsic viscosity of the DNA polymer accounts for the slower coalescence. In theory, the viscosity of condensates should scale as a power of the molecular weight of the polymer[44]. However, under the solvent conditions tested here, and for DNA lengths ¡ 3kbp, the scaling relationship between intrinsic viscosity and DNA length is expected to be near l 2.7kbp DNA required several minutes to complete coalescence (Figure 2D). This greater than 100X increase in the rate of coalescence overshoots our expectations based solely on DNA length changes, demonstrating that HP1*α*-DNA interactions also contribute to the intrinsic viscosity of the condensate. Moreover, condensates formed with 9kbp DNA (~60X larger than 147bp) were unable to complete coalescence within an hour (Figure 2D). And while these condensates do exhibit a slow reduction in perimeter over time, suggesting that coalescence is proceeding locally, at the whole condensate level, the morphology of these condensates remains aspherical. Together these results indicate that within condensates, DNA is constrained by HP1*α* interaction networks leading to novel conformational restrictions and effective polymer interactions. Importantly, the length of heterochromatic domains *in vivo* is typically greater than 10kbp. Therefore, the molecular interactions that occur in condensates formed around longer DNA molecules (9kbp and longer) resulting in non-spherical morphologies may more closely mimic *in vivo* genomic environments.

Overall these experiments suggest that HP1*α* and DNA differentially contribute to bulk droplet properties; the length of DNA and how it interconnects with HP1*α* interaction networks delimits condensate viscosity, while HP1*α* interactions likely define condensate surface tension. This means, that as the DNA length increases, the timescale for global conformational rearrangements of the DNA polymer also increase, while the timescale for rearrangements of HP1*α*-DNA and potentially HP1*α*-HP1*α* interactions are likely to remain fairly constant.

### D. HP1*α* dynamically binds to DNA while simultaneously maintaining stable DNA domains

To further investigate the interplay between these two types of rearrangements (HP1*α*-DNA and HP1*α*-HP1*α* vs. intra-DNA dynamics), we quantified the dynamics of HP1*α* and DNA within condensates We assessed the dynamics of HP1*α* using fluorescence recovery after photobleaching (FRAP). We find that for HP1*α*, despite large differences in droplet morphology, the rate of recovery is unaffected by changes in DNA length after partial photobleaching (Figure 3A-C). This result is consistent with HP1*α*-DNA and potential HP1*α*-HP1*α* interactions remaining unaffected by changes in DNA length. Condensates formed with DNA ranging in length from 147bp to ~50kbp showed recovery of fluorescence with comparable t1/2 values (s) (Figure 3C), which are strikingly similar to recovery rates of HP1*α* measured *in vivo*[16, 17]. Consistently, bleaching of the entire condensate also showed rapid recovery of fluorescence within experimental error of complete recovery (figure supplement 3.1E). These results demonstrate that HP1*α* readily exchanges within condensates, and between condensate and solution populations, without disruption of the condensates. To further test the mobility of HP1*α*, we mixed pre-formed condensates prepared using HP1*α* labeled with either Atto488 (HP1*α*-488) or Atto565 (HP1*α*-565) (Figure 3D, figure supplement 3.2D). Within seconds after mixing, both HP1*α*-488 and HP1*α*-565 were found to have partitioned equally into all droplets (Figure 3D, figure supplement 3.2D). This rapid mixing of fluorescent protein is in full agreement with the FRAP estimates of HP1*α* mobility.

**Figure 3.**
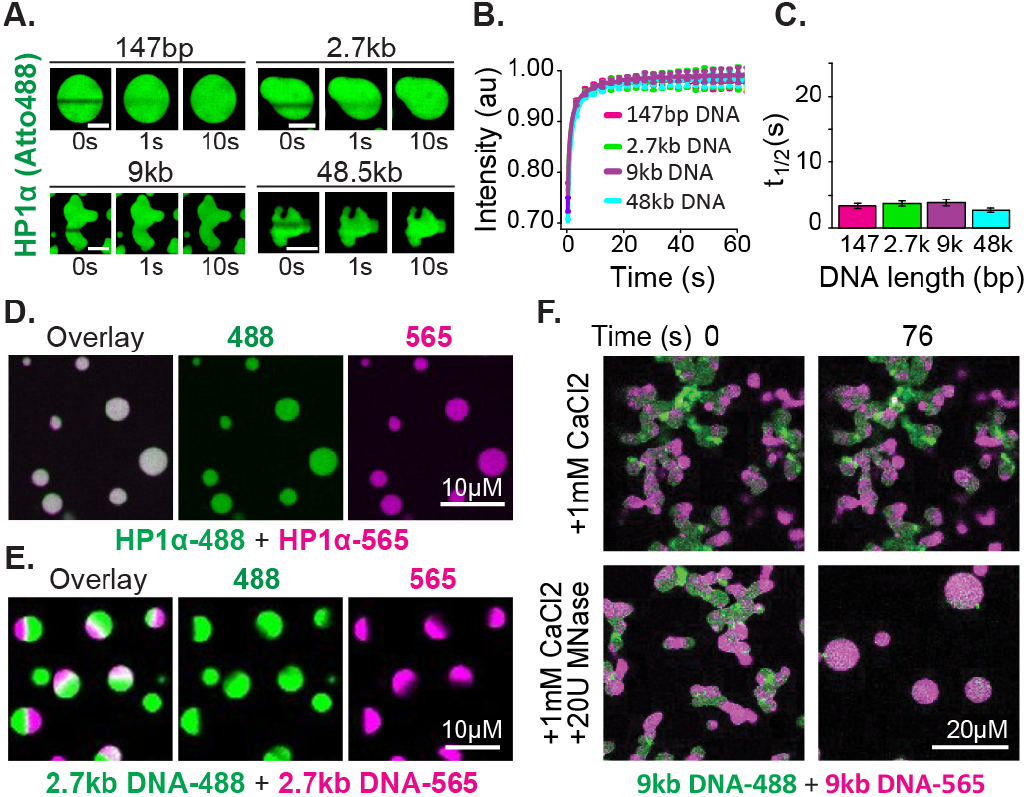
Distinct characteristics of HP1*α* and DNA in condensates. **A.** FRAP of HP1*α* in condensates. Timestamped images from FRAP experiments for fluorescent HP1*α* and four lengths of linear DNA (147bp, 2.7kbp, 9kbp, or 50kbp). Scale bar indicates 5*µ*m. **B.** Recovery of HP1*α* fluorescence intensity and **C.** half-life of HP1*α* recovery plotted for each DNA length tested. **D.** Two color HP1*α* mixing experiments. Condensates formed separately with 2.7kbp unlabeled DNA and either HP1*α*-488 (green) or HP1*α*-565 (magenta) imaged 1.16 minutes after mixing. **E.** Two color DNA mixing experiments. Condensates formed separately with unlabeled HP1*α* and 2.7kbp DNA-488 (green) or 2.7kbp DNA-565 (magenta) imaged 4.4 minutes after mixing. **F.** MNase treatment of condensates. Mixed condensates formed separately with unlabeled HP1*α* and 9kbp DNA-488 (green) or 9kbp DNA-565 (magenta) treated with either 1mM CaCl_2_ or 1mM CaCl_2_ and 20 units of MNase. Images shown for both conditions before and 76 seconds after the treatment.

Next we tested the mobility of the DNA polymer inside condensates. We performed mixing experiments using condensates preformed with HP1*α* and 2.7kbp DNA that was end labeled with either Atto488 (DNA-488) or Atto565 (DNA-565) (Figure 3E, figure supplement 3.2E). The DNA length for these experiments was chosen to be long enough to manifest long polymer effects, but short enough to allow for the completion of coalescence (Figure 2D-E). Contrary to the observations above, DNA does not rapidly mix across condensates after fusion but is instead maintained large and long lived (¿ 1 hour) singlecolor sub-condensate domains (Figure 3E, figure supplement 3.2E). Furthermore, FRAP experiments of HP1*α*DNA condensates labeled with YOYO-1 exhibit recovery rates proportional to DNA length: the longer the DNA, the slower the rate of recovery (figure supplement 3.2AC).

These results confirm substantially different timescales for the mobility of HP1*α* versus DNA, as discussed in the previous section. Further, these results demonstrate that linear DNA as short as 3kbp can be sustained in static compartments, despite prevalent and rapid exchange of HP1*α*. This outcome can arise through either the aforementioned viscosity and conformational constraints inherent to long DNA molecules, and/or through a collective activity of HP1*α* in condensates. To test if DNA viscosity is required for the persistence of sub-condensate DNA domains and non-spherical morphology, we dynamically altered the length of DNA in condensates by the addition of the calcium-dependent non-specific DNA nuclease, micrococcal nuclease. For these experiments, two-color HP1*α*-DNA condensates were formed using 9kbp DNA resulting in diversely shaped condensates with alternating domains of fluorescence (Figure 3F). We expect that if polymer viscosity is required to maintain both the morphology of condensates and the reduced mobility of DNA, dynamically shortening the DNA length should result in both the resumption and completion of coalescence, and uniform mixing of fluorescent signals. Digestion of the DNA reveals this expectation to be accurate, and we observe rapid coalescence and mixing of alternately labeled DNA within condensates (Figure 3F). Importantly, we observe no effects on either phenomenon due to inclusion of calcium alone (Figure 3F).

Overall, these experiments reveal a remarkable character of HP1*α*-DNA condensates—a fast exchanging, liquid pool of HP1*α* can stably trap and organize large DNA molecules into isolated and long-lived domains. Seemingly, HP1*α* accomplishes this feat by increasing the effective viscosity of long DNA molecules to establish and maintain stable condensate structures. This rationale is consistent with our observation that changes to viscosity in HP1*α*-DNA condensates scale more sharply than expected from DNA length considerations alone. We note that the presence of nucleosomes will change the DNA length dependence of viscosity driven effects. However, as we describe in the discussion, these differences will disappear at genomic scales and we expect that HP1 molecules will similarly increase the effective viscosity of chromatin to generate stable chromatin domains.

### E. HP1*α* maintains compacted DNA at relatively high forces

Given the dynamic behavior of HP1*α*, we expected that condensed HP1*α*-DNA structures, although kinetically long-lasting, would be readily dissolved if subjected to biologically relevant forces. To test this hypothesis, we investigated condensate stability against an externally applied force through optical trapping experiments combined with confocal microscopy (Figure 4AB). In these experiments, we performed stretch-relax cycles (SRCs) (figure supplement 4C) by repetitively stretching and relaxing single DNA molecules in presence of HP1*α*. Simultaneously, we measured the force required to extend the DNA to a given length, yielding force-extension curves (Figure 4C, figure supplement 4A). Prior to adding HP1*α*, we first ensured that each tether was composed of a single molecule of DNA and behaved as previously described (Figure 4C)[47]. We then moved the trapped DNA molecule, held at an extension of ~5.5*µ*m, to a chamber containing HP1*α* and observed the formation of compacted HP1*α*-DNA structures analogous to those observed on DNA curtains (Figure 1B, 4B). This initial incubation was sufficiently long to complete condensate formation (30s). Notably, in this assay, compacted DNA structures appear in the center of the DNA molecule rather than at the end, because, with the motion of both ends of the DNA constrained by their attachment to polystyrene beads, the largest DNA chain fluctuations occur in the middle of the molecule.

**Figure 4.**
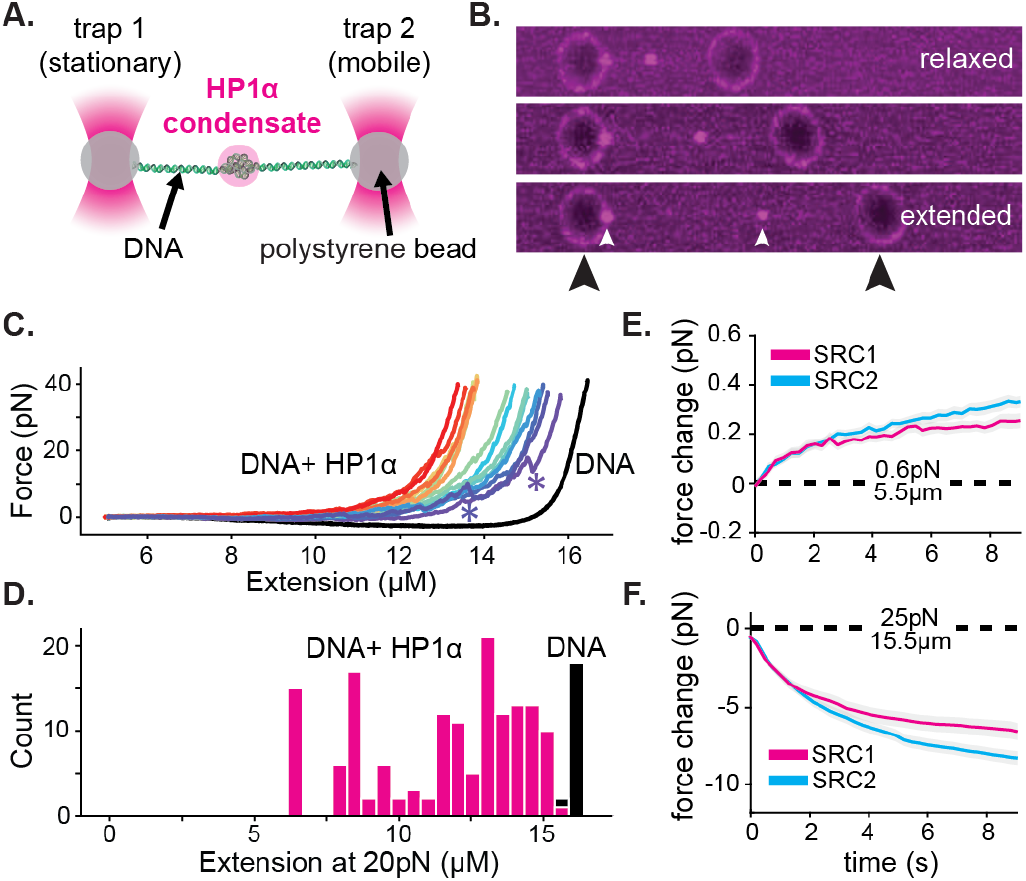
HP1*α*-DNA condensates resist disruptive forces and retain memory of past strain. **A.** Cartoon of optical trap experiments. **B.** Confocal images of relaxed, intermediate, and extended states of DNA (unlabeled) in the presence of HP1*α* (magenta). Black arrowheads indicate trapped beads and white arrowheads indicate HP1*α*-DNA condensates. **C.** Force extension curves for DNA in the absence (black line) or presence of HP1*α* (colored lines). Each trace represents a single stretch-relax cycle (SRC) of the same DNA strand. Traces are colored by pulling order from first extension (violet) to the final extension (red). **\*** indicates rupture event. **D.** Histogram of DNA extension at 20pN in the absence (black) or presence of HP1*α* (magenta). **E.** and **F.** Force change for DNA incubated with HP1*α* in **E.** relaxed or **F.** extended conformation. Shown is the average of the first (magenta) and second (cyan) SRC.

For our initial experiments, DNA tethers bearing internal HP1*α*-DNA condensates were stretched at constant velocity to a final force of 40pN, immediately relaxed, and then stretched again (Figure 4C, figure supplement 4A). We observe a substantial deviation in the force extension curve for DNA in the presence of HP1*α* relative to DNA alone (Figure 4C, figure supplement 4A). We verified that the shift to larger forces for DNA extended in the presence of HP1*α* is not a consequence of radi-ation driven cross-linking (figure supplement 4B). From this measurement, we identify three prominent features of HP1*α*-DNA interactions. First, sequestered DNA domains, measuring on average 10kbp, are able to resist disruption to an instantaneous force of 40pN (Figure 4C,D). However, smaller HP1*α*-DNA structures (1-2kbps) are observed to rupture at lower forces ranging from 5-20pN, suggesting the stability of HP1*α*-compacted DNA scales by size (Figure 4C, see “*”). Second, by integrating the area between the force-extension curves for DNA alone and in the presence of HP1*α*, we estimate that an average energetic barrier of ~1kbT/bp of compacted DNA separates HP1*α*-compacted states of DNA from extended DNA states in the absence of HP1*α* (Figure 4C, figure supplement 4A). Finally, we observed that each successive SRC resulted in more DNA stably sequestered by HP1*α* (Figure 4C). This surprising result shows that, after HP1*α*-DNA condensates are subjected to strain, polymer rearrangements and/or force-dependent selection of HP1*α* binding interactions provide a basis for further stable compaction of DNA by HP1*α*.

Next we asked whether or not HP1*α*-DNA condensates could compact DNA against force or maintain the compacted state when subjected to sustained force by performing consecutive SRCs that included waiting periods after complete relaxation (~5.5*µ*m) and after stretching to 25pN (~15.5*µ*m) (Figure 4E-F, figure supplement 4CE). During the waiting period after relaxation, we observe a steady force increase over time (Figure 4E, figure supplement 4D-E). This result may be the product of either association of HP1*α* molecules from solution and/or rearrangements of DNA and already bound HP1*α*. To test whether low-force DNA compaction required a constant influx of HP1*α* binding, we moved the DNA tether from the chamber containing HP1*α* to a chamber containing only buffer and performed an additional three SRCs (figure supplement 4D). We find that even in the absence of free HP1*α*, the population of already bound HP1*α* molecules is sufficient to induce compaction in the low force regime (~1pN) (figure supplement 4D). Notably, compaction in the absence of free protein can be abrogated by increasing the ionic strength of the buffer (from 70mM to 0.5M KCl) (figure supplement 4E), consistent with salt-induced decompaction observed on DNA curtains[24].

When the DNA is held at a steady extension of 15.5*µ*m following stretching, we observe a drop in measured force over time (Figure 4F, figure supplement 4D-E). This relaxation indicates that HP1*α*-DNA condensates are biased toward disassembly during sustained higher forces. This result is potentially due to force-dependent changes in the kinetics of HP1*α* binding and/or the reduction in DNA strand fluctuations required by HP1*α* to induce compaction. To test whether HP1*α* in solution could affect the stability of the condensate, through a facilitated exchange mechanism[48], we again performed an additional three SRCs in the absence free HP1*α* (figure supplement 4D-E). We find that the disassembly of HP1*α*DNA condensates at higher forces proceeds at the same rate irrespective of the presence of HP1*α* in solution (figure supplement 4D).

Notably, during both waiting periods—before and after stretching—we measure changes in HP1*α*-DNA condensation activity in later SRCs (Figure 4E-F, figure supplement 4D-E). In the relaxed configuration, during low-force compaction, we observe more robust compaction during the second SRC relative to the first (Figure 4E). In comparison, we observe more rapid disassembly while waiting at higher forces during the second SRC (Figure 4F). These strain-induced effects on HP1*α* behavior can have important consequences for how HP1*α*-organized genetic material responds to cellular forces. For example, RNA polymerase ceases to elongate when working against forces as low as 7.5-15pN[49]. Our experiments show that short transient bursts by polymerase are unlikely to disassemble and may even strengthen HP1*α*compacted structures above the force threshold for efficient transcription. However, repeated, sustained efforts by polymerase might be sufficient to relax HP1*α*compacted structures and allow for transcription to proceed.

Moreover, these data suggest that a dynamic network of HP1*α*-DNA and potential HP1*α*-HP1*α* interactions can account for both increased viscosity and stabilization of global condensate structure. In general, we propose that such properties arise from a mean-field activity of an exchanging population of HP1*α* molecules that constrain the DNA at any given time. That is, regardless of the stability of any individual HP1*α* molecule, the average character of the HP1*α*-DNA network is maintained in condensates at a pseudo steady state.

While the measured stability of HP1*α*-DNA condensates is consistent with a role for HP1*α* as a mediator of transcriptional repression, it is hard to reconcile this activity with dynamic chromatin reorganization when cellular cues necessitate the disassembly of heterochromatin. These data also raise the question of which molecular features of HP1*α* allow it to realize its many functions in condensates and on single DNA fibers. Below we first study the molecular features of HP1*α* that drive condensate formation and then address how HP1*α*-DNA condensates may be disassembled.

### F. The hinge domain of HP1*α* is necessary and sufficient for DNA compaction and condensate formation

First, we set out to determine the smallest piece of HP1*α* sufficient for the collective HP1*α* behaviors on DNA we have observed. HP1*α* is comprised of three disordered regions interspaced by two globular domains: a chromodomain (CD) and a chromoshadow domain (CSD) (Figure 5A)[18]. The CD binds to diand trimethylation of lysine 9 on histone 3 (H3K9me) and the CSD mediates HP1 dimerization as well as interactions with other nuclear proteins[7, 21, 50, 51]. The central disordered region, or hinge domain, of HP1*α* mediates DNA binding[25, 52]. Finally, the N-terminal extension (NTE) and the C-terminal extension (CTE) of HP1*α* have been shown to regulate oligomerization of phosphorylated HP1*α*[24]. While all five domains of HP1*α* collaborate to determine *in vivo* localization, cellular localization at heterochromatic sites is completely abolished by mutations to the hinge domain of HP1*α*[16, 53, 54].

**Figure 5.**
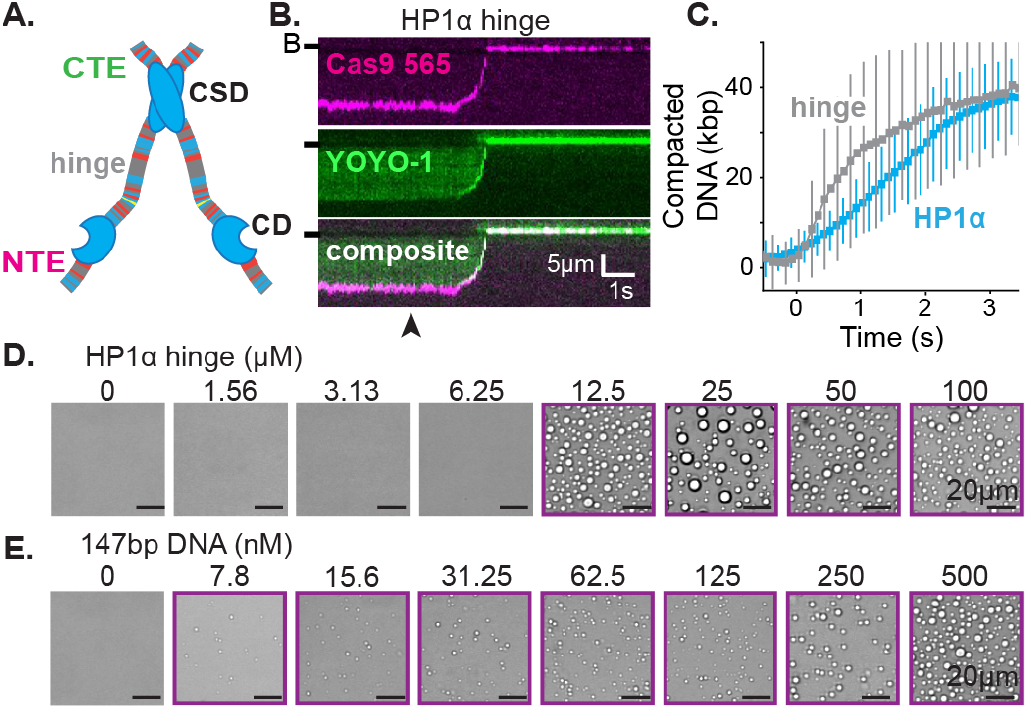
The hinge region of HP1*α* is sufficient for DNA compaction and condensate formation. **A.** Cartoon of HP1*α* with color-coded disordered regions: positive residues (K and R) blue, negative residues (E and D) red, proline yellow, and all other residues grey. Key HP1*α* domains are labeled: chromodomain (CD), chromoshadow domain (CSD), hinge, N-terminal extension (NTE), and C-terminal extension (CTE). **B.** Kymogram of DNA compaction by the hinge domain. DNA is labeled with dCas9 (top) and YOYO-1 (middle), also shown as composite image (bottom). Arrowhead represents estimated time of protein injection. (B-) or (-) specifies location of the barrier. **C.** Average DNA compaction by 5*µ*M HP1*α* and 5*µ*M HP1*α*-hinge. **D.** and **E.** Brightfield images of the HP1*α*-hinge and DNA. **D.** Titration of the HP1*α*-hinge with 500nM 147bp DNA. **E.** Titration of 147bp DNA with 12.5*µ*M HP1*α*-hinge. Purple boxes indicate presence of condensates.

Therefore, we first investigated the activity of the hinge domain isolated from the rest of the protein. Surprisingly, not only is the hinge domain sufficient for DNA compaction (Figure 5B-C, figure supplement 5B-C), but compaction proceeds at twice speed of the full-length protein (Figure 5C, figure supplement 1.1C, 5C). Additionally, the hinge domain is sufficient to induce the formation of condensates with DNA (Figure 5D-E). And, even with short (147bp) DNA oligomers, the critical concentration for condensate formation is reduced by a factor of five relative to full-length HP1*α* (from 50*µ*M to 12.5*µ*M) (Figure 2A, 5D). Surprisingly, the critical concentration is reduced, and DNA compaction increased, even though the valency of the hinge domain alone is ostensibly half that of full-length HP1*α* due to removal of the CSD. Furthermore, and consistent with observations of full-length HP1*α*, condensates formed with the hinge domain exhibit a continuous reduction in size upon lowering DNA concentration, rather than exhibiting a sharp transition between the presence and absence of droplets (Figure 2A, 5E). The strong *in vitro* activity of the hinge domain alone compared to full length HP1*α*, and the requirement of an unperturbed hinge domain for proper function *in vivo*, raise the possibility that the remaining disordered regions of HP1*α* exist to regulate the behavior of the hinge domain.

### G. The disordered extensions of HP1*α* regulate hinge domain activity

Previous work demonstrated that the NTE and CTE of HP1*α* play opposing roles in controlling the phase-separation behavior of phosphorylated HP1*α*[24]. In this context, the CTE acts in an auto-inhibitory role and phosphorylated residues in the NTE promote oligomerization through interactions with the hinge domains in trans (figure supplement 6.1A). We hypothesized that these two disordered terminal extensions may similarly regulate hinge domain activity in the context of DNA-driven HP1*α* phase-separation. To test this possibility, we deleted these extensions of HP1*α*, either separately or in tandem (figure supplement 6.2A).

Removal of the NTE (HP1*α*-ΔNTE) abolished detectable condensate formation with short DNA oligomers and increased the critical concentration for condensate formation with longer DNA (Figure 6A-B). Furthermore, HP1*α*-ΔNTE compacted DNA ~20 times slower than full-length HP1*α* and only managed to compact ~7 kbp of the available ~50kbps (Figure 6C,E, figure supplement 6.2B-C). These results suggest that removal of the NTE lowers the apparent on-rate for DNA binding, and generally raises the free energy of HP1*α*-DNA interactions. However, the compacted structures that do form in our curtain assay persist even after the pulse of HP1*α*-ΔNTE protein exits the flowcell, suggesting that removing the NTE of HP1*α* might not compromise the off-rate of HP1*α* (figure supplement 6.2C). The inhibition of both DNA compaction and condensate formation upon NTE deletion demonstrates that the NTE plays a positive role in each process. Furthermore, these effects are also consistent with the NTE contributing to higher order oligomerization of HP1*α* in the context of DNA binding (see below).

**Figure 6.**
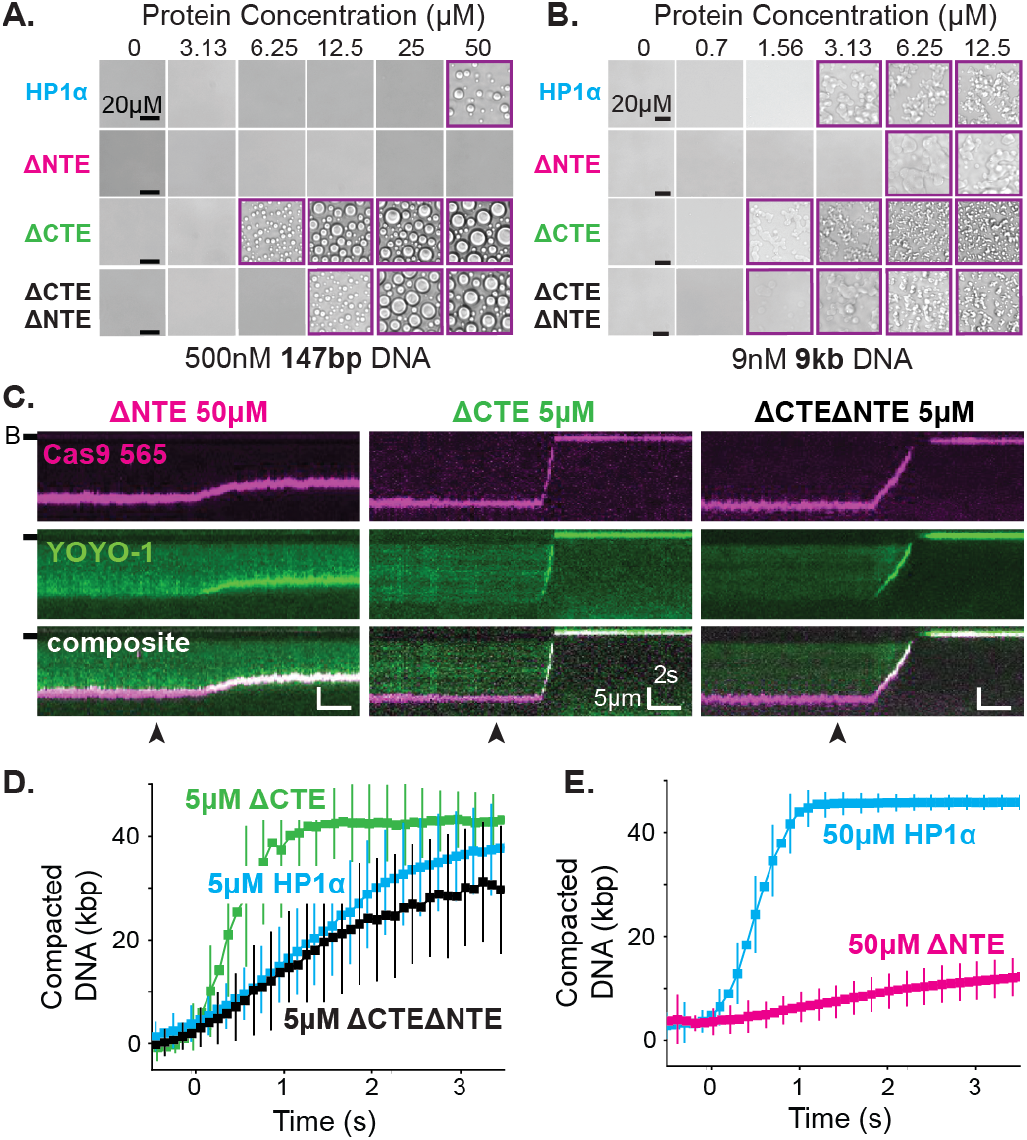
The disordered extensions of HP1*α* regulate DNA compaction and condensate formation. **A.** and **B.** Brightfield images of HP1*α* domain mutants and DNA. **A.** Titration of HP1*α* domain mutants with 500nM 147bp DNA. **B.** Titration of HP1*α* domain mutants with 9nM 9kbp DNA. Purple boxes indicate presence of condensates. **C.** Kymograms of DNA compaction by HP1*α* domain mutants. DNA is labeled with dCas9 (top) and YOYO-1 (middle), also shown as composite image (bottom). Data shown for reactions including 50*µ*M HP1*α*ΔNTE, 5*µ*M HP1*α*ΔCTE, and 5*µ*M HP1*α*ΔNTEΔCTE, respectively. Arrowheads represent estimated time of protein injection. (B-) or (-) specifies location of the barrier. **D.** Average DNA compaction by 5*µ*M HP1*α*, 5*µ*M HP1*α*ΔCTE, and 5*µ*M HP1*α*ΔCTEΔNTE. **E.** Average DNA compaction by 50*µ*M HP1*α* and 50*µ*M HP1*α*ΔNTE.

In contrast, deletion of the CTE (HP1*α*-ΔCTE) decreased the critical concentration for condensation with 147bp DNA oligomers an order of magnitude (Figure 6AB). This result indicates that removal of the CTE lowers the free energy of HP1*α*-DNA condensation. HP1*α*-ΔCTE also compacted DNA three times faster than full-length HP1*α* and almost twice the apparent rate measured for the hinge domain alone (Figure 6C-D, figure supplement 5C, 6.2C). Together with the compaction activity of the hinge and HP1*α*-ΔCTE, these data demonstrate that the CTE negatively regulates the activity of the hinge domain in the context of full-length HP1*α*. This is consistent with previous crosslinking mass-spectrometry studies that indicate the CTE binds to the hinge when not bound to DNA[24].

Finally, when both the NTE and CTE are removed from HP1*α* (HP1*α*-ΔNTEΔCTE), we observe intermediate phenotypes: compaction rates faster than HP1*α*-ΔNTE but slower than HP1*α*-WT, HP1*α*-ΔCTE, or the hinge alone (Figure 6C-E, figure supplement 5C, 6.2C) and a decrease in the critical concentration for HP1*α*DNA condensation, though not to the same extent as HP1*α*-ΔCTE (Figure 6A-B). This result further supports our model of opposing regulation of the hinge domain by the NTE and CTE of HP1*α* in the context of the fulllength protein.

The findings above reveal that the HP1*α* hinge is sufficient for condensate formation with DNA and that its activity is regulated by the CTE and NTE of HP1*α*. In previous sections we have shown that full-length HP1*α* binds to DNA and induces local compaction that nucleates and supports the growth of phase separated domains. Now it is clear that these behaviors are subject to, and resultant of, a complex and coordinated network of interactions between the domains of HP1*α* (xfigure supplement 6.1A). This regulation of activity likely occurs between the disordered domains of individual HP1*α* molecules and also across many HP1*α* molecules throughout HP1*α*-DNA complexes.

### H. Differential droplet formation and DNA compaction by HP1 paralogs

The three human HP1 paralogs, differ significantly in sequence across their unstructured regions (Figure 7AB)[18]. Our results thus far suggest these differences should manifest differential activities with DNA and offer a convenient approach to study the regulation of HP1*α*’s hinge domain by the NTE and CTE. First, we tested each paralog’s ability to compact DNA (Figure 7C-E). We find that HP1*β* displays a substantially reduced rate of DNA compaction relative to HP1*α* (Figure 7C,E). This indicates a relative deficiency in the apparent interaction strength between HP1*β* and DNA. Indeed HP1*β*’s compaction activity is more comparable to that of HP1*α*-ΔNTE (figure supplement 6.2C, 7A-C). Yet, despite slower compaction, HP1*β* continues to sustain compacted DNA even after the bulk of the injected pulse of HP1*β* has passed through the flowcell (Figure 7E, figure supplement 7A-C). This suggests a lower bound for HP1*β*’s off-rate from compacted DNA on the order of minutes. In comparison, HP1*γ* compacts DNA more rapidly than HP1*β*, though HP1*γ* does not achieve the rapid compaction rates of HP1*α* (Figure 7D-E, figure supplement 7A-C). Moreover, HP1*γ* rapidly disassembles as the concentration of free HP1*γ* in the flowcell begins to decline (Figure 7E, figure supplement 7A-C). We propose that this effect is the result of HP1*γ* having a faster off-rate from DNA relative to HP1*α* or HP1*β*. Importantly, these experiments suggest that genomic regions organized by HP1*α* and HP1*β* would require less protein for maintenance and be more resistant to disruption relative to domains organized by HP1*γ*.

**Figure 7.**
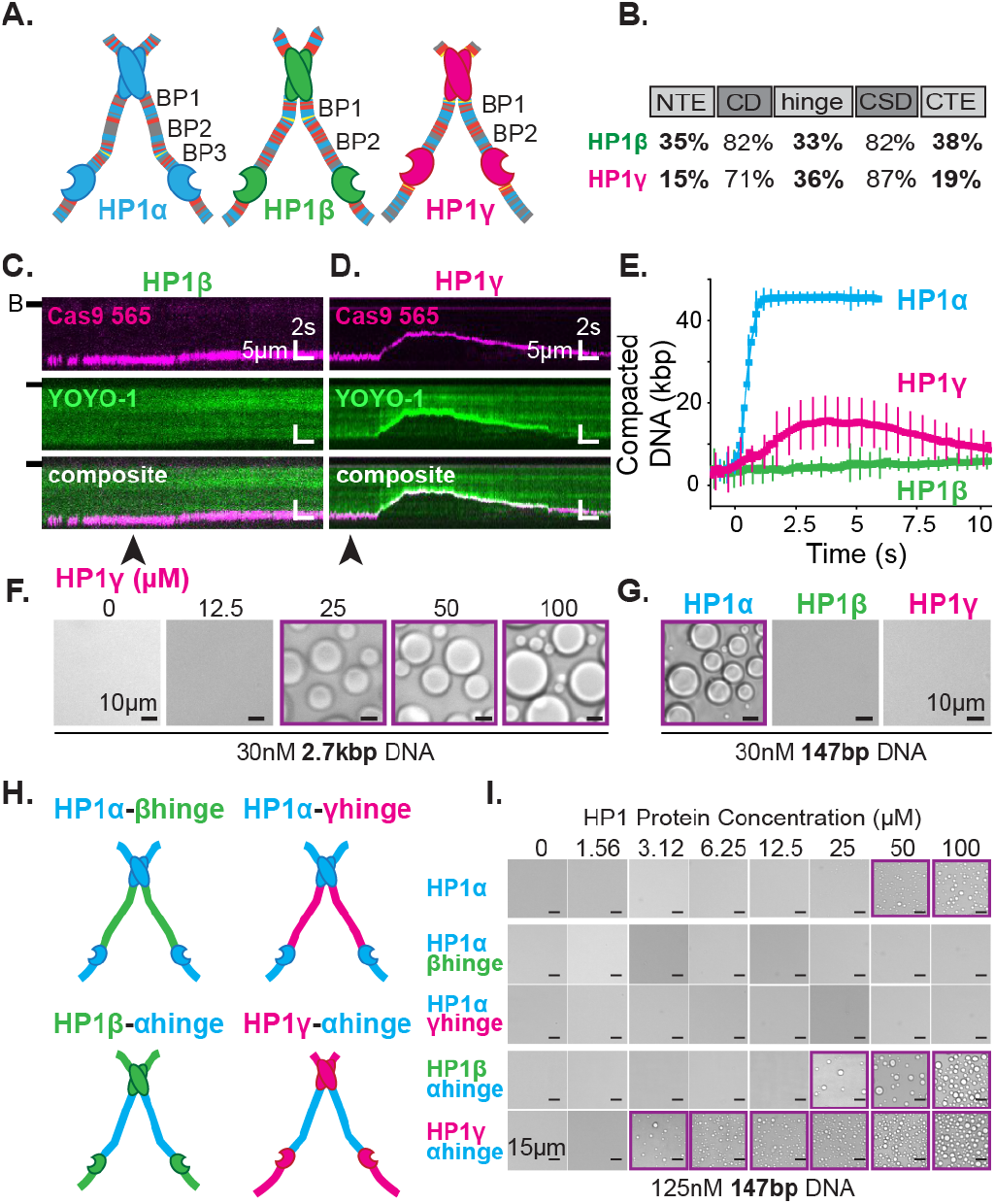
DNA compaction and condensate formation activity of HP1*β* and HP1*γ*. **A.** Cartoons of the three paralogs of human HP1 with color-coded disordered residues: positive residues (K and R) blue, negative residues (E and D) red, proline yellow, and all other residues grey. Basic patches (BP) for each homolog are labeled. **B.** Comparison of amino acid homology between HP1*α* and HP1*β* or HP1*γ*. **C.** and **D.** Kymograms of DNA compaction by **C.** HP1*β* and **D.** HP1*γ*. DNA is labeled with dCas9 (top) and YOYO-1 (middle), also shown as composite image (bottom). Arrowheads represent time of protein injection. (B-) or (-) specifies location of the barrier. **E.** Average DNA compaction by 50*µ*M HP1*α*, HP1*β*, and HP1*γ*. **F.** Brightfield images of HP1*γ* and 2.7kbp DNA. **G.** Brightfield images of 100*µ*M HP1*α*, HP1*β*, or HP1*γ* and 147bp DNA. H.Cartoon of HP1 hinge domain swaps. I.Brightfield images of HP1 domain swap mutants and 147bp DNA. Purple boxes indicate presence of condensates.

Next we tested our interpretation of compaction experiments by assessing the relative capacity of each HP1 paralog to drive condensate formation with DNA (Figure 7F-G). We predicted that due to its decreased compaction rate, HP1*β* would struggle to form condensates with DNA. However, if any condensates form, we would predict that those HP1*β*-DNA structures would be stable. On the contrary, we expect HP1*γ* will readily condense into liquid domains with DNA but require a higher concentration to maintain condensation relative to HP1*α*, due to the apparent increase in reversibility of compaction on curtains (Figure 7D-E, figure supplement 7A-C). We find that HP1*γ* does form condensates with 3kbp DNA, though the critical HP1*γ* concentration required to induce droplet formation is, in fact, higher than that for HP1*α* (Figure 2E, 7F). Moreover, HP1*γ* does not form condensates with 147bp DNA, under conditions where HP1*α* continues to drive DNA condensation (Figure 7G). These results are consistent with a lower DNA binding affinity and higher off-rate for HP1*γ*. In contrast, we find that HP1*β* does not induce droplet formation regardless of the length of co-incubated DNA (Figure 7G, data for longer DNA not shown). This result mirrors the attenuated condensate forming activity of HP1*α*-ΔNTE and is consistent with lower DNA compaction rates and a lower DNA binding affinity. Furthermore, HP1*β* demonstrates that the ability to induce and maintain stable DNA compaction in it of itself is not definitive of condensate formation.

### I. The disordered regions of HP1 paralogs drive differential DNA compaction and condensate formation activity

The above results uncovered substantial differences in the abilities of HP1*β* and HP1*γ* to compact and form condensates with DNA as compared to HP1*α*. We presumed these differences in activity are due to differences in their respective disordered domains. Specifically, we expect the disparities across paralogs in their hinge domain, which for HP1*α* is sufficient for DNA compaction and condensation (Figure 5B-E), to be the strongest predictor of activity. To directly determine the differences in activity due to individual hinge domains, we replaced the hinge domain of HP1*α* with the corresponding hinge domains from either HP1*β* or HP1*γ*, respectively (HP1*α*-*β*hinge and HP1*α*-*γ*hinge) (Figure 7H). We find that both mutants fail to produce condensates in the presence of DNA (Figure 7I), demonstrating that, within the context of full-length HP1*α* and the HP1 paralogs, the HP1*α*’s hinge domain is necessary for droplet formation. While it might have been expected for HP1*α*-*γ*hinge to exhibit some condensate formation activity, it is worth noting that HP1*γ* lacks any appreciable CTE, and its NTE is remarkably different than that of HP1*α* (Figure 7A-B). Therefore, in its native context, the hinge domain of HP1*γ* likely does not have to navigate autoregulation in order to promote productive HP1*γ*-DNA interactions.

We then performed compensatory swaps of the hinge domain of HP1*α* into HP1*β* (HP1*β*-*α*hinge) and HP1*γ* (HP1*γ*-*α*hinge) (Figure 7H). We find both of these mutants now readily form condensates with DNA, demonstrating the HP1*α* hinge is also sufficient for phase separation in the context of the other HP1 paralogs (Figure 7I). Intriguingly, the critical concentration for condensate formation was decidedly lower for both *α*-hinge mutants than for HP1*α*; two-times lower for HP1*β*-*α*hinge and ten-times lower for HP1*γ*-*α*hinge (Figure 7I). These results indicate that the HP1*α* hinge is more active outside of its native context where it is free from the inhibitory effect of its CTE.

The HP1 paralogs are often found in overlapping genomic regions in cells. Given the differential activities of the paralogs, we next asked now mixed populations might manifest distinct properties in condensates by performing droplet assays in the presence of paralog competitors. When HP1*β* or HP1*γ* were premixed with HP1*α* and added to DNA to assess condensate formation, both HP1*β* and HP1*γ* inhibited droplet formation in a concentration dependent fashion (Figure 8A). Notably, these experiments were performed with 147bp DNA, which when incubated with HP1*γ*, does not induce condensate formation (Figure 7G). Interestingly, when introduced to preformed HP1*α*-DNA condensates, HP1*β* is capable of invading and subsequently dissolving condensates at a rate proportional to HP1*α* exchange (Figure 8B). In contrast, HP1*γ* does not destabilize, but rather enriches in the pre-formed HP1*α*-DNA condensates (Figure 8C). These results may simply be a reflection of binding site competition. However, the HP1 paralogs have been suggested to heterodimerize, so it is attractive to hypothesize that heterodimers between HP1*α* and HP1*β* or HP1*γ* have lower DNA binding affinity or disrupted regulatory interactions such that condensate formation is inhibited. Furthermore, while it is difficult to account for the differences in pre-formed condensate disruption by HP1*β* and HP1*γ* with a simple steric occlusion model, differences in heterodimerization activity and/or activity of heterodimers provide an acceptable rationale.

**Figure 8.**
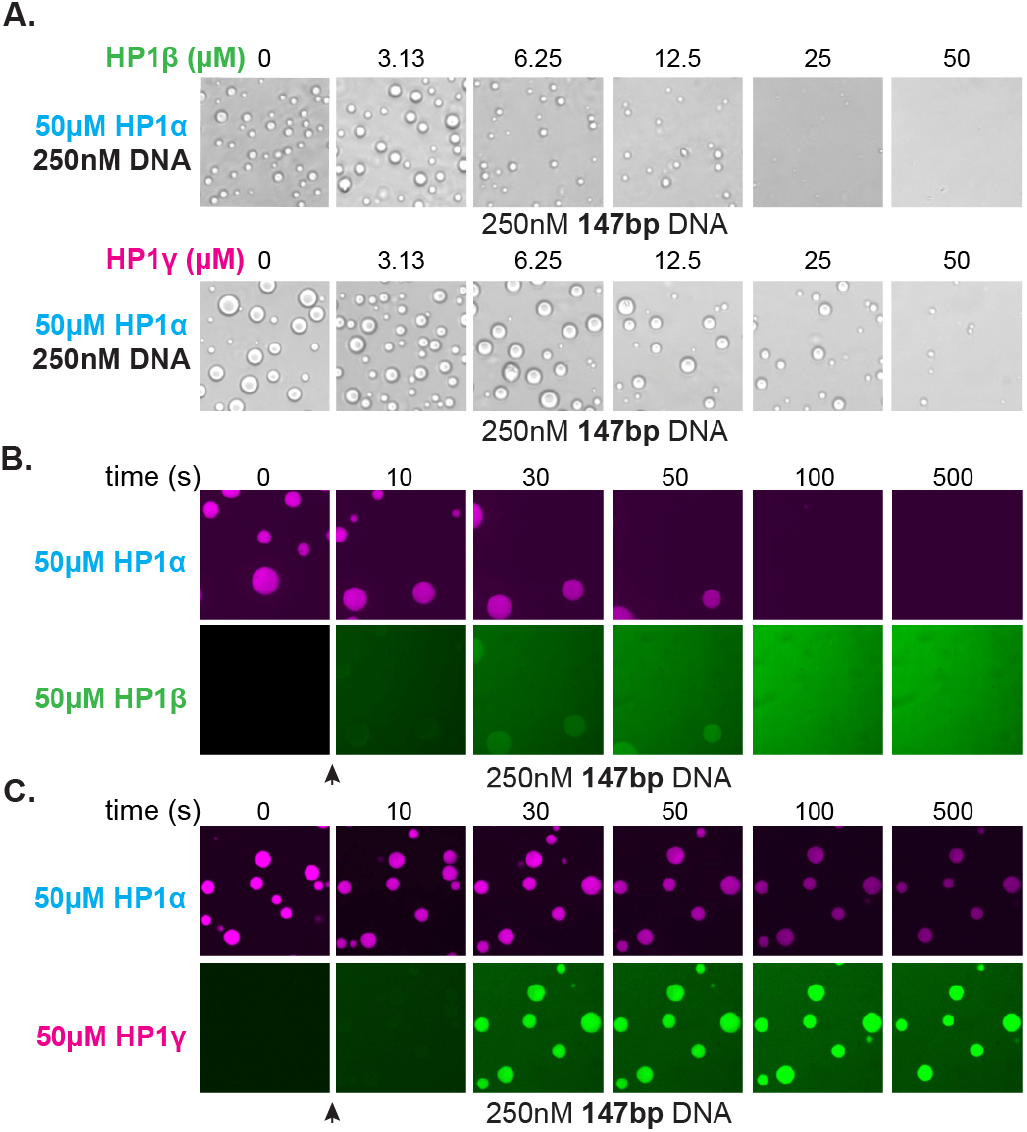
Effect of HP1*β* and HP1*γ* on HP1*α*-DNA condensate formation and stability. **A.** Brightfield images of DNA and pre-incubated mixtures of HP1*α* and HP1*β* (top) or HP1*α* and HP1*γ* (bottom). **B.** and **C.** Confocal images showing a time course of HP1*α* condensates after injection of **B.** HP1*β* or **C.** HP1*γ*.

Together, these results suggest inter-paralog competition as a possible mechanism of cellular regulation of HP1-mediated chromatin domains. Moreover, these experiments demonstrate the critical advantage of biological organization by liquid condensates—competition can be fast. Fast competition means that, regardless of domain stability, when the molecular environment changes, condensates can respond to those changes at the rate at which the organizing material exchanges. For condensation of DNA by HP1*α*, this means that even in the context of highly viscous, tangled DNA and large networks of protein-protein and protein-DNA interactions that resist mechanical disruption at steady state, domains can easily be disassembled in seconds due to the rapid exchange rate of individual HP1*α* molecules.

## III. DISCUSSION

Heterochromatin serves to organize large regions of the eukaryotic genome into domains that are positionally stable yet can be disassembled in response to cell cycle and developmental cues[11–16]. Previous work on HP1-mediated heterochromatin uncovered several key biophysical properties of HP1 proteins such as the ability to form oligomers and to form liquid-like phase-separated condensates with DNA and chromatin[21, 23, 24, 28–30]. A closer examination of these properties can help discern their cellular influence and ultimate role in regulation of heterochromatin states in cells. However, a major challenge in such an endeavor has been connecting the actions of individual HP1 molecules on DNA to the collective phenotype of a heterochromatin domain. Here, we have used a series of complementary assays that allow us to measure the mesoscale behavior of human HP1 proteins and interpret that behavior in terms of single molecule activity. Our findings indicate at least three regulatable steps by which HP1*α* organizes and compacts DNA (Figure 9A-C): (1) Local assembly of HP1*α* along DNA prior to DNA condensation; (2) initiation of DNA compaction through capturing of proximal DNA fluctuations via HP1*α*-DNA and HP1*α*-HP1*α* interactions to form a proto-condensate, and (3) progression of DNA compaction through inclusion of uncompacted DNA into the growing condensate via HP1*α*-DNA and HP1*α*-HP1*α* interactions. We further find that the polymer behavior of DNA, together with the ability of HP1*α* molecules to make multivalent interactions with rapid on/off kinetics, results in stable mesoscale structures that resist mechanical forces but are subject to competition (Figure 9C-D). Finally, comparison of the behavior of HP1*α* with that of HP1*β* and HP1*γ* uncovers new biophysical differences between the three paralogs. Below we discuss the mechanistic and biological implications of our findings in the context of previous observations.

**Figure 9.**
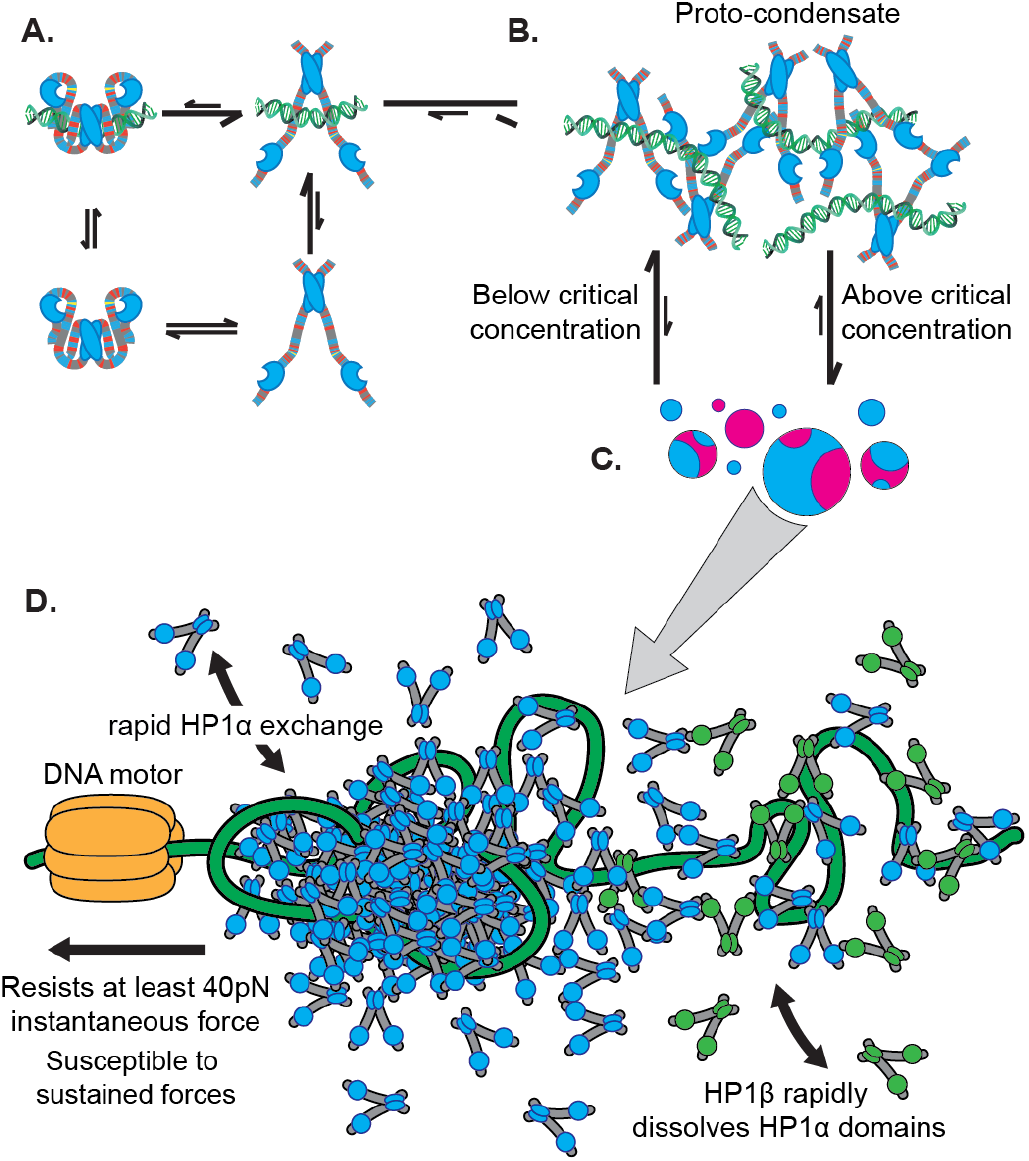
Microscopic to macroscopic activity of HP1*α*. **A.** At the microscopic scale, interactions between the terminal extensions and hinge domain toggles HP1*α* between autoinhibited and active states. DNA biases HP1*α* to the active state. **B.** At the intermediate scale, HP1*α* and DNA cluster into proto-condensates. **C.** If HP1*α* is present above the critical concentration, proto-condensates aggregate into large macroscopic droplets characterized by liquid behavior of HP1*α* and static DNA held in sub-condensate domains. **D.** At genomic loci, HP1*α* condensates are remodeled by forces, resisting and strengthening in response to instantaneous forces, but relaxing and weakening in response to sustained forces. HP1*α* domains are also subject to disruption and reinforcement from HP1-interacting proteins like HP1*β*.

### A. Implications for regulation of heterochromatin assembly and spreading

The framework presented above has implications for understanding how heterochromatin domains grow through incorporation of additional regions of the genome. Specifically, factors that lower the affinity of HP1*α* for DNA, or potentially HP1*α*’s affinity for itself, will result in reduced formation of compacted DNA and a heightened sensitivity to disruption. Regions of DNA that are low affinity binding sites for HP1*α* will also resist incorporation into compacted domains and can potentially act as insulating sites against HP1*α* activity.

Furthermore, in our experiments, we find that longer DNA promotes the formation of HP1-DNA condensates (Figure 2A,E, 6A-B, 7F-G). This observation is consistent with longer DNA, with higher valency, increasing the local concentration of proto-condensates. Therefore, restricting the continuity of HP1*α* binding sites *in vivo* would also be predicted to inhibit growth of heterochromatin domains. An obvious way to interrupt continuous stretches of DNA is by the presence of nucleosomes. Indeed ¿70% of mammalian genomes are occupied by nucleosomes[55]. The traditional view is that H3K9me3 containing nucleosomes act as platform for HP1 interactions that impart preference for heterochromatin versus euchromatin[9]. In this context, it is tempting to speculate that histone modifications act to restore HP1 binding sites interrupted by the nucleosome core, thereby promoting HP1 assembly and specificity. At the same time, the presence of nucleosomes would also regulate the architecture of HP1 assembly. Consistent with such a possibility, HP1 proteins from S. pombe have been shown to bridge across and deform H3K9me3 nucleosomes[28]. Furthermore, interactions made by HP1 proteins with the histone octamer and H3 tail may serve additional roles in regulating the stability of the condensate.

The internal regulatory network of interactions across the hinge, NTE, and CTE regions of HP1*α* will also influence assembly on chromatin *in vivo*. Our results imply that protein binding or post-translational modification of the NTE and CTE could have large effects on the ability of HP1*α* to condense chromatin. For example, proteins that bind to the CTE may induce HP1*α* to behave more like HP1*α*-ΔCTE promoting condensation (Figure 6A-D, figure supplement 6.2B-C). Indeed, a large number of nuclear proteins bind HP1*α* in close proximity to the CTE, including two proteins shown to modulate HP1*α* phase separation *in vitro*, Lamin-B receptor and Shugoshin[24, 51]. Alternatively, modifications may provide the basis for new interactions as seen when the N-terminal of HP1*α* is phosphorylated[24]. Importantly, because HP1*α* concentrations in the cell are similar to the lower limit for condensate formation that we observe *in vitro* (Figure 2E), the assembly of HP1*α* is well-poised to be influenced by molecular interactions and modifications[56].

### B. Implications for the versatility of heterochromatin function

A major function of heterochromatin is the compartmentalization of the genome[3]. In this context, our results indicate a dominant role for the DNA polymer in regulating its own compartmentalization. In condensates, we find that ~3 kbp pieces of DNA are fixed in place on the order of an hour, while HP1*α* molecules can diffuse on the order of seconds (Figure 3, figure supplement 3.1, 3.2). Our results imply that such behavior arises from two sources: the intrinsic viscosity of DNA due to its polymer properties, and the mean activity of rapidly rearranging HP1*α* molecules, which creates an average protein-DNA network equivalent to a set of static interactions. As a result, when two condensates fuse, the HP1*α* molecules rapidly exchange between the two condensates while the DNA from each condensate remains trapped in separate territories (Figure 3D-E, figure supplement 3.2D-E).

Inclusion of nucleosomes substantially increases the persistence length and linear density of DNA while potentially decreasing the number of HP1 binding sites[57]. Since HP1 interactions also contribute to viscosity, any effects from a reduction in HP1 binding sites would be balanced by the increased rigidity of the chromatin polymer. Additionally, in the context of chromatin, HP1 molecules can use additional domains, such as the CD and the CSD, to further constrain chromatin through interactions with H3K9me modifications and the histone core, respectively. In all these considerations it is important to note that the length effects due to the large sizes of chromatin domains in the nucleus would overshadow differences between the viscosities of chromatin versus DNA. Thus, we propose that the meso-scale behaviors observed in the context of DNA will be recapitulated in the context of chromatin, but with additional regulatable steps.

From a charge passivation perspective, the ability of HP1*α* to condense DNA bears similarities to counterion mediated condensation of DNA by ions such as spermidine[58]. Interestingly, spermidine mediated DNA condensates dissolve upon application of ~1pN force requiring only ~0.1kT/bp of work in contrast to the ¿1kbT/bp required to disassemble HP1*α*-DNA condensates(Figure 4C)[35]. Some these differences may arise from the specific DNA binding properties of the hinge region as opposed to those of spermidine. However, the key difference allowing for the higher stabilization of compacted DNA achieved by HP1*α* is the formation of a network of HP1*α*-HP1*α* interactions in addition to HP1*α*DNA interactions.

Importantly, we find that HP1*α*-DNA condensates are able to resist disruption by instantaneous forces of at least 40pN (Figure 4C, figure supplement 4A). Furthermore, we find that transient forces increase the ability of condensates to resist subsequent disruptions (Figure 4C,E, figure supplement 4D-E). The high resistance to force, as well as the conversion to a more stable state upon application of transient external force, provides a biophysical explanation for how heterochromatin can confer mechanical stability in two contexts: to the nucleus when the nuclear membrane is subjected to mechano-chemical signaling events, and to centromeres when they are subjected to forces of chromosome segregation[3, 10]. However, we also show that sustained high forces provoke the relaxation of condensates and sensitize condensates for subsequent disruptions (Figure 4F, figure supplement 4D-E). These effects highlight the ability of HP1-mediated heterochromatin to be shaped by cellular forces.

Our results further explain how mechanically stable and long-lived domains are dissolved in response to cellular cues. We find that, even while the global character of HP1-DNA condensates is fixed, the constituent HP1 molecules are highly dynamic (Figure 3, figure supplement 3.1, 3.2)[23]. This dynamism allows for rapid competition and interference, and, because the organizing network of HP1-DNA interactions is built from weak transient encounters, results in swift disassembly of structures and dispersal of material (Figure 8B). More generally, because condensates often rely on the integrated weak interactions of large populations to build cellular structures, they also present low energetic barriers to competition. Condensates thus present unique advantages in the context of cellular organization. It is, however, worth noting that competition need not be direct and the chemical environment in condensates can also restrict competitor access to internal structures. The general organizational principles that we have uncovered here can be applied in many biological contexts but seem most readily applicable to the unique functions and constraints shared by genome organizing proteins.

### C. Implications for paralogs and evolution

In addition to HP1*α*, there are two other paralogs of HP1 in humans, HP1*β* and HP1*γ*. Despite sharing similar domain architecture and conservation of sequence, these paralogs of HP1 differentially localize in the cell and perform individual functions (Figure 7A-B)[59, 60]. Importantly, each paralog also performs distinctly in our two assays. We find that HP1*β* binds to DNA at a lower rate than HP1*α*, leading to reduced DNA compaction activity; yet, the compaction by HP1*β* is relatively stable (Figure 7C,E, figure supplement 2.2D, 7A-C). Additionally, we find that HP1*β* is unable to produce observable condensates with DNA (Figure 7G). This may be because HP1*β* is deficient in modes of DNA binding, is unable to engage in protein-protein interactions beyond dimerization, its central disordered region is ill adapted to condensation, or any combination therein. Notably, when HP1*β* also makes nucleosomal contacts, it can compact chromatin leading to condensation[30, 61]. Additionally, HP1*β* is a particularly effective competitor for HP1*α in vitro*, suggesting that HP1*β* interactions may be adapted for tempering HP1*α*-organized chromatin or to serve a role in establishing chromatin boundaries (Figure 8A-B). Furthermore, we find HP1*γ* binds to DNA at a much faster rate than HP1*β*, but HP1*γ*-DNA condensates also rapidly disassemble in the absence of excess free protein, resulting in rapid compaction followed by rapid decompaction on DNA curtains (Figure 7D-E, figure supplement 2.2D, 7A-C). Yet, HP1*γ* is able to induce condensate formation with DNA, though at a higher protein concentration than HP1*α* (Figure 7F-G). Notably, in certain cells, HP1*γ* is the most diffuse HP1 paralog, often not exhibiting localization at all, which might be the result of the high instability we observe[59, 60]. The higher critical concentration for HP1*γ*-DNA condensation reflects a higher setpoint for regulation in comparison to HP1*α*, meaning HP1*γ* will require a larger cellular investment in protein levels to induce condensation. It is also possible that higher order chromatin organization by HP1*γ* may be at cross purposes with the known role of HP1*γ* in promoting transcription elongation[62].

The three human HP1 paralogs are the result of past gene duplications, and while they have faithfully conserved their chromoand chromoshadow domains, their disordered regions have diverged completely (Figure 7B)[63, 64]. It is possible that each paralog achieves specificity in biological function through their disordered regions. We demonstrate this possibility by exchanging disordered domains among the paralogs, converting HP1*β* and HP1*γ* into robust agents of DNA condensation (Figure 7H-I). These experiments also reveal the evolutionary potential of the modular HP1 domain architecture. For example, it’s easy to imagine the effect that inserting a sequence with variable condensing ability into HP1*α* would have on the genome and heterochromatin stability. Indeed, the molecular diversity of HP1 proteins across eukaryotes suggests that evolution has already taken advantage of HP1 architecture[64].

### D. Implications for the diversity of biological liquid-like phase-separation phenomenon

In addition to heterochromatin, phase-separation phenomena have been observed in many different biological contexts, including the nucleolus, P bodies, and P granules[65–69]. Recent studies have also found evidence for phase-separation behavior in the context of transcription and DNA repair[70, 71]. The increasing number of observations of phase-separation in biological systems has created an apparent need to define the criteria for liquid phase-separated condensates[32]. Some commonly proposed criteria for liquid-like phase-separation behavior are, (i) a boundary that confines the mobility of phase-separating molecules, (ii) concentration buffering, and (iii) differential viscosities inside versus outside of condensates[31, 32, 65]. Some of these criteria are based on assumptions of a homogenous solute (condensed phase) and solvent (surrounding phase). However *in vivo*, both the solute and solvent are heterogenous. Below we discuss how this, and other considerations make the above criteria limiting in the context of biologically meaningful condensates.

#### 1. Boundaries

It has been proposed that phase-separated condensates will have boundaries that promote preferential movement of phase-separating molecules inside the condensate as opposed to movement of molecules across the boundary. To measure such a property, recent studies have assessed the permeability of condensate boundaries to the entry and exit of GFP-HP1*α* molecules using FRAP, and shown that entry of GFP-HP1*α* molecules from the outside the condensate occurs at least as fast as the rate of internal mixing of GFP-HP1*α* molecules[31]. However, for rapidly moving molecules, like HP1*α*, and small domain sizes, like chromocenters, differences in recovery rates due to internal mixing versus exchange across the boundary will be near resolution limits. Furthermore, *in vitro*, some liquid condensates have been demonstrated to have surprisingly low density and high permeability[72]. In these condensates it is possible to decouple the movement of molecules within condensates from their mesoscale droplet properties. Indeed, our results show that HP1*α* molecules *in vitro* can mix both within and across condensates at comparable rates (Figure 3A, figure supplement 3A-C, figure supplement 3.1). Furthermore, our measured rates of HP1*α* exchange (seconds) in condensates are likely to make measurements of differential dynamics within versus without condensates difficult to distinguish. Importantly, the results from our FRAP studies of HP1*α*-DNA condensates *in vitro* reveal exchange rates that are similar to prior FRAP studies on heterochromatin puncta in cells[16, 17, 31]. Overall, our results demonstrate that some categories of condensates can display mesoscale liquid-like characteristics even while the motion of molecules within condensates and across their boundaries are similar.

#### 2. Concentration buffering

Concentration buffering refers to a phenomenon where increasing the total concentration of a condensing protein, like HP1*α*, does not change the concentration inside relative to outside of condensates[65]. Instead, the volume of condensates increases. In opposition, recent studies have shown that increasing the cellular concentrations of HP1*α* by overexpression does not result in an increase in the size of heterochromatin puncta but instead increases the concentration of HP1*α* inside the puncta[31].

Interestingly, our *in vitro* data is consistent with some expectations of concentration buffering of HP1*α* above the critical concentration. Specifically, we show that the size of HP1*α*-DNA condensates grows with the addition of either DNA or HP1*α* (Figure 2C). However, it is important to note that partitioning of material into condensed and soluble phases is also defined by the energetics of the molecular interactions in either compartment. In the *in vitro* context, concentration buffering in HP1*α*-DNA condensates would be achieved if there is only competition for binding interactions between HP1*α* molecules, and if the chemical environment inside of the condensates does not vary as a function of this competition. However, it is also possible that additional HP1-HP1 interactions may change the concentration buffering behavior of HP1*α*DNA condensates. Further, in cells, heterochromatin is expected to contain other DNA-binding factors in addition to HP1*α*. Therefore, it is possible that increasing concentrations of HP1*α* simply compete off other component molecules within heterochromatin puncta. Critically, it has recently been shown that this assumption of concentration buffering also fails to describe the concentration dependence of protein inclusion in the nucleolus, perhaps the best-defined phase-separated organelle in the cell[33].

#### 3. Differential viscosities

*in vitro* studies using nucleolar components have shown that some nucleolar proteins such as NPM1 substantially increase the bulk viscosity of condensates compared to water[67]. This result has led others to define liquidlike phase-separated condensates by whether or not they exhibit increased viscosity relative to the outside dilute phase[31]. However, there are limitations to using differential viscosities a defining feature of liquid-like phase-separation. For example, a recent study used fluorescence correlation microscopy of GFP-HP1*α* to measure the viscosity of GFP-HP1*α* in chromocenters in cells. Based on similar rotational diffusion of GFP-HP1*α* inside and outside of heterochromatin puncta, the study concluded that the HP1*α* in chromocenters does not experience a higher viscosity relative to elsewhere in the nucleus and therefore do not conform to the definitions of LLPS[31]. However, as opposed to micro-rheology experiments which measure bulk viscosities[67], rotational diffusion reports on a convoluted energetic landscape defined by a multitude of interactions made by the diffusing molecules. Specifically, in this case, the conclusion that HP1*α* does not engage in LLPS *in vivo* relies on the validity of the assumption that there is a distinct separation between the strength and abundance of molecular interactions inside relative to outside condensates. For simple *in vitro* systems diffusion of condensing material will almost always be slower inside condensates rather than outside, as the outside represents a dilute molecular environment. However, in the nucleus, there is no analog of the “dilute phase” as the majority of the nucleus is crowded with diverse molecules. Thus, it is possible for HP1*α* to make interactions both within and outside of condensed heterochromatin *in vivo* that results in comparable mobilities relative to a dilute solution.

### E. Conclusion

The discussion above about the complexity of the cellular environment relative to *in vitro* experiments raises the general question of how *in vitro* demonstrations of liquid-like phase-separation can be used to derive biologically meaningful insights. At a foundational level, quantitative *in vitro* experiments are essential to detail the properties of the biological condensates, as we have done for the HP1-proteins. The complexities of cellular contexts can then be layered on, and tested, in a systematic manner. Critically, determination that the simplest assumptions of LLPS behavior derived from *in vitro* studies are not upheld *in vivo* is extremely valuable in identifying the effects of such additional complexity[31]. But, if molecules exhibit liquid-like phase separation activity *in vitro*, the interactions that produce that behavior do not vanish in the cell. Rather they are integrated into the complex network of cellular interactions that spans all of the molecules in the cell. And importantly, the interactions that give rise to macroscopic LLPS *in vitro* are present even among sparingly few molecules, regardless of whether they manifest across scales into large liquid domains. Sometimes those interactions will be obscured by cellular activity, but in other contexts those same interactions may be at the forefront biological activity. *in vitro* studies are therefore essential to provide a framework to test and interpret the relative behaviors of phase-separating components in cells. Overall, our findings here underscore that, as new activities of biological condensates continue to be discovered it is important to characterize the biophysical nature of these condensates and the biologically relevant properties that they enable.

## IV. MATERIALS AND METHODS

### A. Protein Purification

General method: Rosetta competent cells (Millipore Sigma 70954) transformed with expression vectors for 6x-HIS tagged HP1 proteins were grown at 37^*◦*^C to an OD600 of 1.0-1.4 in 1 liter of 2x LB supplemented with 25*µ*g/mL chloramphenicol and 50*µ*g/mL carbenicillin. HP1 protein expression was induced by the addition of 0.3mM isopropy-*β*D-thiogalactopyranoside (IPTG). Cells were then grown for an additional 3 hours at 37^*◦*^C, before pelleting at 4,000xg for 30 minutes. Cell pellets were then resuspended in 30mL Lysis Buffer (20mM HEPES pH7.5, 300mM NaCl, 10% glycerol, 7.5mM Imidazole) supplemented with protease inhibitors (1mM phenylmethanesulfonyl fluoride (Millipore Sigma 78830), 1*µ*g/mL pepstatin A (Millipore Sigma P5318), 2*µ*g/mL aprotinin (Millipore Sigma A1153), and 3*µ*g/mL leupeptin (Millipore Sigma L2884). Cells were then lysed using a C3 Emulsiflex (ATA Scientific). Lysate was clarified by centrifugation at 25,000xg for 30 minutes. The supernatant was then added to 1mL of Talon cobalt resin (Takara 635652) and incubated with rotation for 1 hour at 4^*◦*^C. The resin-lysate mixture was then added to a gravity column and washed with 50mL of Lysis Buffer. Protein was then eluted in 10mL of elution buffer (20mM HEPES pH 7.5, 150mM KCl, 400mM Imidazole). Then, TEV protease was added to cleave off the 6x-HIS tag and the protein mixture was dialyzed overnight in TEV cleavage buffer (20mM HEPES pH 7.5, 150mM KCl, 3mM DTT) at 4^*◦*^C. The cleaved protein was then further purified by isoelectric focusing using a Mono-Q 4.6/100 PE column (GE Healthcare discontinued) and eluted by salt gradient from 150mM to 800mM KCl over 16 column volumes in buffer containing 20mM HEPES pH 7.5 and 1mM DTT. Protein containing fractions were collected and concentrated in a 10K spin concentrator (Amicon Z740171) to 500*µ*L and then loaded onto a Superdex-75 Increase (GE Healthcare 29148721) sizing column in size exclusion chromatography (SEC) buffer (20mM HEPES pH7.5, 200mM KCl, 1mM DTT, 10% glycerol). Protein containing fractions were again collected and concentrated to 500*µ*M in a 10K spin concentrator. Finally, aliquots were flash frozen in liquid nitrogen and stored at −80^*◦*^C.

HP1*α*, HP1*β*, and HP1*γ* were all purified as described above. For the terminal extension deletes (HP1*α*ΔNTE, HP1*α*ΔCTE, and HP1*α*ΔNTEΔCTE) minor changes to the ionic strength of buffers were made. Specifically, each protein was dialyzed into a low salt TEV protease buffer (20mM HEPES pH 7.5, 75mM KCl, and 3mM DTT) in the overnight cleavage step. Additionally, the salt gradient used in isoelectric focusing ranged from 75mM to 800mM KCl. The rest of the protocol followed as written above.

The HP1*α* hinge was purified as written until the overnight TEV cleavage step. After which, the protein was loaded onto a Hi-Trap SP HP column (GE Healthcare 17115201) and eluted in a salt gradient from 150mM to 800mM KCl over 16 column volumes in buffer containing 20mM HEPES and 1mM DTT. Protein containing fractions were collected and concentrated in a 10K spin concentrator to 500*µ*L and then loaded onto a Superdex30 10/300 increase (GE Healthcare 29219757) sizing column in size exclusion chromatography (SEC) buffer. Protein containing fractions were then collected and concentrated to 500*µ*M in a 10K spin concentrator. Finally, aliquots were flash frozen in liquid nitrogen and stored at −80^*◦*^C.

### B. Protein labelling

Proteins constructs for fluorescent labelling were modified to contain a C-terminal GSKCK tag and to substitute native reactive cysteines to serine residues (HP1*α*C133S and HP1*γ*-C176S). For labeling, HP1 proteins were dialyzed overnight into SEC buffer with 1mM TCEP substituted for DTT. Protein was then mixed at a 1:1 molar ratio with either maleimide Atto488 or maleimide Atto565 (Millipore Sigma 28562, 18507). The reaction was immediately quenched after mixing by addition of 10x molar excess of 2-mercaptoethanol. Labeled protein was then separated from free dye over a Hi-Trap desalting column (GE Healthcare 17-1408-01) in SEC buffer. Labeled protein was then flash frozen in liquid nitrogen and stored at −80^*◦*^C.

### C. DNA purification

Plasmids containing DNA used in this study were amplified in DH5*α* cells (ThermoFisher 18265017) grown in TB. Plasmids were purified using a Qiagen Plasmid Giga kit (Qiagen 12191). Plasmids containing the “601” DNA sequence were digested with EcoRV (NEB R0195S) and the 147bp fragments were then isolated from the plasmid backbone by PAGE purification. Briefly, DNA were loaded into a 6% acrylamide gel and run at 100mV for ~2 hours in 1xTBE. The desired 147bp DNA band was cut out of the gel and soaked in TE (10mM Tris-HCL pH 7.5, 1mM ETDA) buffer overnight. The supernatant was then filtered, and DNA isolated by two sequential ethanol precipitations. The 2.7kbp DNA (Puc19) was linearized by HindIII (NEB R0104S) digestion and purified by two sequential ethanol precipitations. The 9kbp DNA (pBH4-SNF2h[73]) was linearized by BamHI (NEB R0136S) digestion and purified by two sequential ethanol precipitations. DNA from bacteriophage *λ* (*λ*-DNA) (NEB N3011S) used in phasing and curtains experiments was prepared by heating to 60^*◦*^C to release base pairing of the cohesive ends in the presence of complementary 12bp primers as previously described [34, 74]. For curtain experiments, the primer targeted to the 3’ overhang of *λ*-DNA was modified to include a 5’ biotin. *λ*-DNA and primers were then allowed to slowly cool to room temperature and then incubated overnight with T4 DNA ligase (NEB M0202S). The *λ*-DNA was then precipitated in 30% PEG(MW 8000) and10mM MgCl_2_ to remove excess primers and washed 3 times in 70% ethanol before resuspension and storage in TE.

### D. DNA labelling

DNA was end-labeled with fluorescent dUTPs as follows. 50*µ*g linear 2.7kbp and 9kbp plasmids were incubated with 12.5 units of Klenow 3’ 5’ exofragment (NEB M0212S), 33 *µ*M dATP, dCTP, dGTP (Allstar scientific 471-5DN), and either 33*µ*M of either ChromaTide Alexa Fluor 568-5-dUTP (ThermoFischer Scientific C11399) or ChromaTide Alexa Fluor4885-dUTP (ThermoFischer Scientific C11397) in 1xT4 DNA ligase buffer (NEB B0202S) at room temperature overnight. Fluorescently labeled DNA was then purified by ethanol precipitation, resuspended in 1xTE, and dialyzed overnight in 1xTE to remove any residual nucleotides.

DNA was biotinylated by performing fill-in reactions with 5 units of Klenow 3’ 5’ exofragment and 0.8mM dTTP, 0.8mM dGTP, 3.2*µ*M bio-dCTP, 8*µ*M bio-dATP (NEB N0446S, Thermo Fisher 19518018, R0081). The reaction was incubated at room temperature overnight and then DNA were purified by ethanol precipitation. Purified DNA were then resuspended in 1xTE to a working concentration of 4mg/mL.

### E. Curtain Assays

DNA curtain experiments were prepared and executed as described elsewhere [34]. Briefly, UV lithography was used to pattern chromium onto a quartz microscope slide, which was then assembled into a flowcell (Figure 1A). A lipid bilayer was established within the flowcell by injecting a lipids mix containing 400*µ*g/mL DOPC, 40*µ*g/mL PEG-2000 DOPE, and 20*µ*g/mL biotinylated DOPE (Avanti Polar Lipids 850375, 880130, and 870273) diluted in lipids buffer (10mM Tris pH 7.5, 100mM NaCl). Streptavidin, diluted in BSA buffer (20mM HEPES pH7.5, 70mM KCl, 20*µ*g/mL BSA, and 1mM DTT), was then injected into the flowcell at a concentration of 30*µ*g/mL. Biotinylated DNA from bacteriophage *λ*, prepared as described above, was then injected into the flowcell and anchored to the bilayer via a biotin-streptavidin linkage. Buffer flow was then used to align the DNA at the nanofabricated barriers and maintain the curtain in an extended conformation during experiments. End-labeling of DNA was accomplished using dCas9 molecules. Specifically, dCas9 (IDT 1081066), Alt-R CRISPR-Cas9 tracrRNA (IDT 1072532), and an AltR CRISPR-Cas9 crRNA targeting bacteriophage *λ* at position 47,752 (AUCUGCUGAUGAUCCCUCCG) were purchased from IDT (Integrated DNA Technologies). Guide RNAs were generated by mixing 10*µ*M crRNA and 10*µ*M tracrRNA in in Nuclease-Free Duplex Buffer (IDT 11050112), heating to 95^*◦*^C for 5 min and then slowly cooling to room temperature. Guide RNAs were then aliquoted and stored at −20^*◦*^C. To prepare Cas9 RNPs for labeling, 200nM of dCas9 mixed with 1*µ*M of guide RNA in dCas9 Hybridization Buffer (30mM HEPES pH 7.5 and 150mM KCl) and incubated for 10 minutes at room temperature. Next, 166nM of the dCas9-RNA mixture was incubated with 0.08mg/mL of 6x-His Tag Antibody conjugated with Alexa Fluor 555 (Invitrogen MA1-135A555) on ice for 10 minutes. Labeled RNPs were then diluted in BSA buffer and injected into the flowcell at a final concentration of 4nM. Labeled dCas9 were allowed to incubate with DNA in the flowcell for 10 minutes before being washed out using imaging buffer (BSA Buffer supplemented with an oxygen scavenging system consisting of 50nM protocatechuate 3,4-dioxygenase (Fisher Scientific ICN15197505) and 31*µ*M protocatechuic acid (Abcam ab142937)). Experiments where DNA are labeled, imaging buffer included 20pM YOYO-1 (Thermo Fisher Y3601).

For compaction experiments, HP1 proteins were diluted to the stated concentration in imaging buffer and injected into the flowcell at a rate of 0.7mL/min. The volume of protein injected was decided based on protein concentration: for experiments with 50*µ*M protein, 100*µ*L was injected, for 5*µ*M protein, 200*µ*L was injected, and for 500nM protein, 400*µ*L was injected. For experiments utilizing fluorescent HP1, labeled protein was included at the following amounts: 200nM HP1*α*-488 was included in the injection 50*µ*M HP1*α*, 400nM HP1*β*-488 was included in the injection of 50*µ*M HP1*β*, and 400nM HP1*γ*-488 was included in the injection of 50*µ*M HP1*γ*. After each experiment, HP1 was removed by washing 0.5M KCl, and replicates performed. Data was analyzed as described below.

### F. Tracking fluorescence during compaction

We measure the fluorescence intensity of both HP1*α*488 and YOYO-1 during DNA compaction. For this analysis, individual ROIs of DNA compaction are segmented manually (Figure 1G). Data were collected for the average and total fluorescence intensity, mean position, and area of both the compacted and uncompacted segments of the DNA.

#### 1. Conservation of YOYO-1 fluorescence

To evaluate our analysis of fluorescence signals due to protein binding, we first tested whether the fluorescence signal from YOYO-1 is conserved across the compacted and uncompacted segments of the DNA during compaction. Assuming YOYO-1 binding is at an equi librium and uniformly distributed across the DNA, we expect the following to be true: i) the total intensity of the uncompacted segment, *I_u_*, is at a maximum before any compaction begins. ii) the total intensity of the compacted segment, *I_c_*, is at maximum at the end of compaction, and iii) max *I_u_* = max *I_c_*. For the analysis we do the following: (1) at each time frame *I_u_* and *I_c_* are measured. (2) *I_u_* is then fit to a line. (3) The value from the linear fit of *I_u_* is subtracted from *I_c_*. And finally (4) *I_c_ I_u_* is normalized by dividing through by max *I_c_*. The final value, *I_c_ I_u_/* max *I_c_* follows our expectations spanning [−1, 1] and crossing zero at the midpoint in the compaction process (figure supplement 1.1A).

#### 2. Association rates of fluorescent HP1α

To investigate the association of HP1*α* during compaction we measure the increase in fluorescent signal along the DNA in both uncompacted and compacted DNA regions. We find on the uncompacted segment of the DNA, the average fluorescent HP1*α* signal per DNA-containing pixel, *ρ*(*t*), increases linearly (Figure 1I). For the compacted regions of DNA, the rate of HP1*α* fluorescence increase is complicated by compaction—the fluorescence can increase both from the association of HP1*α* from solution and from incorporation of HP1*α*-bound DNA into the growing compacted segment.

We first consider how the increase in fluorescence would look if HP1*α* binds to both uncompacted and compacted segments identically. Then the fluorescence intensity of the compacted segment, *I_c_*(*t*), would be equal to:

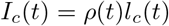

where *l_c_* is the length of compacted DNA and *ρ*(*t*) is the fluorescence density at time *t* from above. Alternatively, if upon condensation, DNA becomes unable to incorporate more HP1*α* from solution, the rate of fluorescence increase over a time will lag behind the first scenario by a factor of ≈ 0.5. This means that if we normalize our measurement of fluorescence intensity, by dividing through by *ρ*(*t*)*l_c_*(*t*), the resulting trend should have no slope in the case of equal binding to both the compacted and uncompacted segments of DNA. Alternatively, a negative slope would indicate that binding to the compacted DNA is impaired relative to uncompacted DNA and a positive slope suggests that binding to the compacted segment is enhanced relative to the uncompacted DNA (Figure 1J-L).

### G. Tracking DNA compaction

To track the length of the DNA we first use an automated program to locate DNA within our images (figure supplement 1.1B). This method is described here Once we have the DNA identified, we make kymograms of each individual DNA strand (figure supplement 1.1C). To make the kymograms, we average over the three rows of pixels local to each DNA strand and stack up the average slice across the frames of the video. Then the kymograms are smoothed using a Gaussian filter. After smoothing, we then take the derivative of the image (figure supplement 1.1C). The derivative is generally at its lowest value at the edge of the DNA where the intensity drops off to background levels, and we set the minimum value, by row, of the derivative filtered image as the end position of the DNA (note the directionality of the derivative is top to bottom) (figure supplement 1.1C). In addition to smoothing the image prior to taking the derivative, we perform two added filtering steps on the data. First, we discount pixels near the edge of the image from the analysis. This is both because often these pixels are added to the kymograms as padding for output and because we know the end of the DNA is not located off image. The second filter is to account for the fact that we expect a relatively smooth trajectory of the DNA end during compaction. To select for this, we take the positions of the DNA end as determined by the minimum of the derivative and apply a Savitzky-Golay filter (SciPy.org). Then, measurements that are more than a few pixels off this smoothed line are discarded. The general analysis pipeline is automated; however, all fits are manually inspected, and fits deemed to be poor due are removed.

### H. HP1*α* binding site size

The end to end distance of an HP1*α* dimer in the closed conformation is 12.9nm. The end to end distance of a phosphorylated HP1*α* dimer phosphorylated in the open conformation is 22.2nm[24]. Assuming 0.34nm/bp, we estimate the minimal binding unit of a HP1*α* dimer in the open conformation is *∼*65bp.

### I. Condensate assays

HP1 condensates were imaged using microscopy grade 384 well plates (Sigma-Aldrich M4437). Prior to use, individual wells were washed with 100*µ*L of 2% Hellmanex (Sigma-Aldrich Z805939) for 1 hour. Then wells were rinsed 3 times with water and 0.5M NaOH was added to each well for 30 minutes before again rinsing 3 times with water. Next, 100*µ*L of 20mg/mL PEG-silane MW-5000 (Laysan Bio MPEG-SIL-5000), dissolved in 95% ethanol, was pipetted into each well and left overnight at 4^*◦*^C protected from light. Next, wells were rinsed 3 times with water and 100mg/mL BSA (Fisher Scientific BP1600) was pipetted into each well and allowed to incubate for 30 minutes. Finally, wells were rinsed 3 times with water and 3 times with 1x phasing buffer (20mM HEPES pH 7.5, 70mM KCl, and 1mM DTT) was added to each well. Care was taken to maintain 10*µ*L of volume at the bottom of the well in all steps to prevent drying of the PEG Silane coating of the bottom of the well. In preparation of experiments, HP1 proteins and DNA substrates were dialyzed overnight into 1x phasing buffer. Then, Protein and DNA were added to a 1.5mL microcentrifuge tube at 1.5x of the final concentration stated in results. Excess phasing buffer was removed from cleaned wells and exactly 10*µ*L of 1x phasing buffer was added to the bottom of the well. Then 20*µ*L of the protein-DNA solution was then added to the well, resulting in a 30*µ*L solution of DNA and protein at the concentrations reported in the results section.

To generate the phase diagram for HP1*α* (Figure 2A), determine condensate radius (Figure 2B, figure supplement 2.2) and for general condensate assays in Figure 2D-E, figure supplement 2.1B-C, Figure 5D-E, Figure 6 A-B, Figure 7F-G,I, and Figure 8A, condensates were visualized by brightfield microscopy at 20X magnification. Condensates were prepared as described above and allowed to incubate for 1 hour at room temperature before imaging. However, for droplet coalescence assays (Figure 2D), droplets were visualized immediately after the reactions were added to the well.

The assays in Figure 3A-F, Figure 8B-C, figure supplement 3.1, and figure supplement 3.2 were imaged by spinning disk confocal microscopy at 100x Magnification. For the mixing assays in Figure 3D and figure supplement 3.2D, 100*µ*M HP1*α* was mixed with 50ng/*µ*L 2.7kbp DNA in 1x phasing buffer for five minutes in two separate reactions with an additional 200nM HP1*α*-488 or 200nM HP1*α*-565 added to each reaction. Then, a single-color reaction was added to a well, briefly imaged, followed by addition of the remaining reaction. The DNA mixing experiments in Figure 3E and figure supplement 3.2E were performed identically to above, except the reactions were prepared using either 50ng/*µ*L 2.7kbp-488 or 50ng/*µ*L 2.7kbp-565 and unlabeled protein.

For the MNase assays in Figure 3F, condensates were formed by incubating 50*µ*M HP1*α* and either 12.5ng/*µ*L 9kbp-488 or 12.5ng/*µ*L 9kbp-565 for 5 minutes. Then individual reactions were mixed and incubated at room temperature for one hour prior to imaging. MNase digestion was initiated by the addition of 1mM CaCl_2_ and 20U MNase (NEB M0247S) and mock reactions were initiated by addition of 1mM CaCl_2_ alone.

For the competition experiments in Figure 8A, HP1*α* was first mixed with either HP1*β* or HP1*γ* to the stated final concentrations. This solution was then added to 147bp DNA (250nM final concentration) and allowed to incubate for 1 hour at room temperature prior to imaging. For the competition experiments in Figure 8BC, condensates were formed with 50*µ*M HP1*α*, 200nM HP1*α*-565, and 250nM 147bp DNA and incubated for 1 hour at room temperature before briefly imaging. Then, either HP1*β*-488 or HP1*β*-488 was added to the reaction to final concentrations of 50*µ*M unlabeled protein and 200nM fluorescent protein.

### J. Droplet segmentation analysis

Many images of HP1-DNA condensates were collected by brightfield microscopy. Segmenting these droplets presented multiple challenges. For example, the rings of high and low intensity at the edges of the droplets and the fact that the intensity inside droplets is almost the same as background intensity. These factors made analysis with basic threshold segmentation difficult. To overcome these difficulties, we created a custom approach utilizing edge detection and several filters (figure supplement 2.1B). We first high pass filter the image in Fourier space. Then we detect the edges of condensates with a Canny edge detector (scikit-image.org). Canny edge detection applies a Gaussian filter to smooth the image before taking the gradient. We found that larger condensates were detected more readily when larger values for the variance of the Gaussian filter were used and smaller condensates when smaller values were used. To implement adaptive smoothing, we calculated the edges across a range of sigma values before combining the segments into a single detected image. This method introduced a significant amount of noise. To remove this noise, we utilized two thresholds: one for condensate area (condensates must be larger than 3 pixels) and the other for condensate eccentricity (condensates must have eccentricity at or less than 0.94). We segmented at least five separate images for each DNA and protein concentration tested and collected the radius of each detected condensate (Figure 2B-C). Then we determined the complementary cumulative distribution (CCD) for condensate radius at each condition (figure supplement 2.2). Confidence intervals for each CCD were determined by the Bootstrap method (figure supplement 2.2). Finally, each curve was integrated to determine the expectation value of the radius for each condition (Figure 2B-C).

### K. FRAP assays

For FRAP experiments, condensates were formed with 100*µ*M HP1*α*, 250nM HP1*α*-488, and 50ng/*µ*L of either linear 147bp, 2.7kbp, 9kbp, or 48.5kbp DNA (see above, DNA purification). Samples were then imaged at room temperature (and 5% CO_2_ for line FRAP experiments). For each photobleaching experiment, automatic focus was activated, pixel binning was set at 2×2, and exposure time was set to 300ms. For line FRAP experiments, a 3×512 pixel rectangle was irradiated with 7mW power at 476nm (Integrated Laser Engine, Andor) for 300ms between the 25th and 26th acquired frame. For the whole droplet FRAP experiments, a custom rectangle surrounding a single condensate was irradiated with 7mW power at 476nm for 1.5s between the 10th and 11th acquired frame. Recovery times to half max (*t*_1*/*2_) were calculated using a biexponential fit.

#### 1. Line FRAP analysis

Line FRAP analysis was performed with a custom Rscript. Unbleached condensates, used for normalization, were segmented by threshold. The ROI of bleached regions of condensates (FRAP ROI) was user-defined during imaging. The intensity of the bleached and unbleached condensates as well as background were measured over time. First, the background was subtracted from the FRAP ROI and the unbleached droplets. Then, the FRAP ROI was normalized via the following equation:

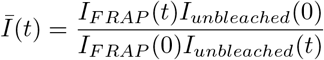

The normalized intensity was then plotted versus time (Figure 3B, figure supplement 3.1B) and fit to a biexponential fit to determine *t*_1*/*2_ values (Figure 3C, figure supplement 3.1C).

#### 2. Whole condensate FRAP analysis

Condensates were formed with 100*µ*M HP1*α*, 250nM HP1*α*-488, and 50ng/*µ*L 2.7kbp DNA and imaged as described. A square ROI incorporating an entire droplet was photobleached and recovery visualized over ten minutes (figure supplement 3.2A). Unbleached condensates, used for normalization, were segmented by threshold. Due to diffusion and, potentially, the chemical environment of condensates, HP1*α* fluorescence decays differently inside of droplets relative to background. Therefore, we only use the signal from the fluorescent HP1*α* within droplets to correct for fluorescence recovery. Additionally, intensity values near the boundary of droplets were omitted from the analysis due to intensity fluctuations resulting from droplet motion. Furthermore, droplets local to the bleached condensate are affected by the bleach strike and are removed from the analysis. Then, we fit the time-dependent decay of condensate fluorescence to a biexponential decay equation (figure supplement 3.2B-C).

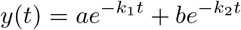

We would then normalize the intensity of the bleached condensate by dividing through by the average decay of unbleached droplets from this equation. However, the intensity of the fluorescent HP1*α* also decays differently depending on its location within the field of view due to non-homogenous illumination of the sample (figure supplement 3.2D). We therefore scale the decay rates of the unbleached droplets in the following way to correct for spatial variation:

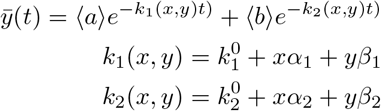

where *α* and *β* and 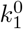 and 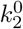 the slopes and intercepts from a linear regression of decay rate versus position in the image, *a* and *b* are the average population factors, and 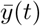 is the adjusted intensity signal. Next, we use the average corrected rates from all of the unbleached condensates to normalize the intensity versus time for all the unbleached condensates. We then use the normalized unbleached intensity versus time to visualize the expected spread of the data, which we use as a visual measure of error (figure supplement 3.2E). Finally, we plot the normalized intensity of the bleached condensate against this unbleached distribution to visualize the extent of fluorescence recovery (figure supplement 3.2E)

### L. Optical trapping experiments

Optical trapping experiments were performed on a Lumicks C-Trap G2 system (Lumicks) or a custom-built dual trap. Trapping experiments were performed in specialized flowcells with separate laminar flow channels. For each experiment, two streptavidin coated polystyrene beads (Spherotec SVP-40-5), diluted to 2.2nM in HP1 buffer (20mM HEPES pH 7.5, 70mM KOAc, 0.2mg/mL BSA, 1mM DTT), were captured. Then, the two beads were moved into a channel containing biotinylated *λ*DNA diluted to 0.5*µ*g/mL in HP1 buffer. Then, using an automated “tether-finder” routine, a single strand of DNA was tethered between two beads. Each DNA strand was stretched at a rate of 0.1*µ*m per second to a maximal force of 40pN in the buffer-only channel two separate times to measure the force extension curve without HP1 present. Next, trapped DNA molecules were moved to a flow channel containing 10*µ*M HP1*α* and 400nM HP1*α*565 and incubated at 5*µ*m extension for 30 seconds. We then perform stretch-relax cycles (SRC) either with or without waiting periods in the extended or relaxed configurations (figure supplement 4C).

For SRCs with no waiting periods (Figure 4C, figure supplement 4A), we performed fifteen SRCs to a maximal force of 40pN in HP1 buffer with 10*µ*M HP1*α* and 400nM HP1*α*-565. For SRCs with waiting periods, we performed three consecutive SRCs to a maximal force of 25pN in HP1 buffer with 10*µ*M HP1*α* and no additional fluorescent protein. We then moved the DNA tether into a channel containing either HP1 buffer or HP1 buffer supplemented with 500mM KCl and performed three additional SRCs (figure supplement 4D-E).

### M. Anisotropy

Prior to anisotropy experiments, HP1*α*, HP1*β*, and HP1*γ* were dialyzed overnight into binding buffer (20mM HEPES pH 7.5, 70mM KCl, and 1mM DTT) at 4^*◦*^C. 60bp DNA oligos containing a 5’-FAM modification (supplementary table) were purchased from IDT (Integrated DNA technologies) and diluted to a final concentration of 10nM in reactions. Binding reactions were then performed in binding buffer supplemented with 0.1mg/mL BSA and variable amounts of HP1 proteins as indicated. Reactions were incubated for 30 minutes at room temperature in Corning Low Volume 384 well plates (Corning LCS3821) then measurements were performed on an Analyst HT (Molecular Devices). Data from three independent HP1 titrations were fit to a one site binding curve and presented with standard errors.

## ACKNOWLEDGMENTS

Special thanks to Lumicks for their generous donation of a C-trap system to the scientific community at the Marine Biological Laboratories during the summer of 2019. Special thanks as well to the Biomolecular Nanotechnology Center at the University of California, Berkeley for their assistance in manufacturing flowcells for DNA curtains. M.M.K. was supported by the Discovery Fellows Program at UCSF and NCI grants F31CA243360 and F99CA245719. R.R. was support from the NOMIS foundation, Rostock, Germany. B.H. acknowledges support though NIH R21 GM129652, R01 CA231300 and R01 GM131641. B.H. is also a Chan Zuckerberg Biohub Investigator. S.W.G. was supported by the DFG (SPP 1782, GSC 97, GR 3271/2, GR 3271/3, GR 3271/4) and the European Research Council (grant 742712).

G.J.N supported through NIH grant GM127020. Support to S.R. through the UCSF Program for Breakthrough Biomedical Research (PBBR), Sandler Foundation, and Whitman Foundation at the Marine Biological Laboratories.

## AUTHOR CONTRIBUTIONS

M.M.K., G.J.N., and S.R. identified and developed the core questions. M.M.K. performed the bulk of the experiments, assisted in data analysis of all experiments and assisted in writing the manuscript. D.B. and B.H. assisted with FRAP measurements and analysis. L.D.B., R.R., and S.W.G. assisted with optical tweezer measurements and analysis. H.K manufactured the microfluidic flowcells used in curtains experiments. C.R.C. assisted with protein purification and experiments involving domain swaps. G.J.N. and S.R. oversaw the research and co-wrote the manuscript with input from all authors.

---

**TABLE I:**
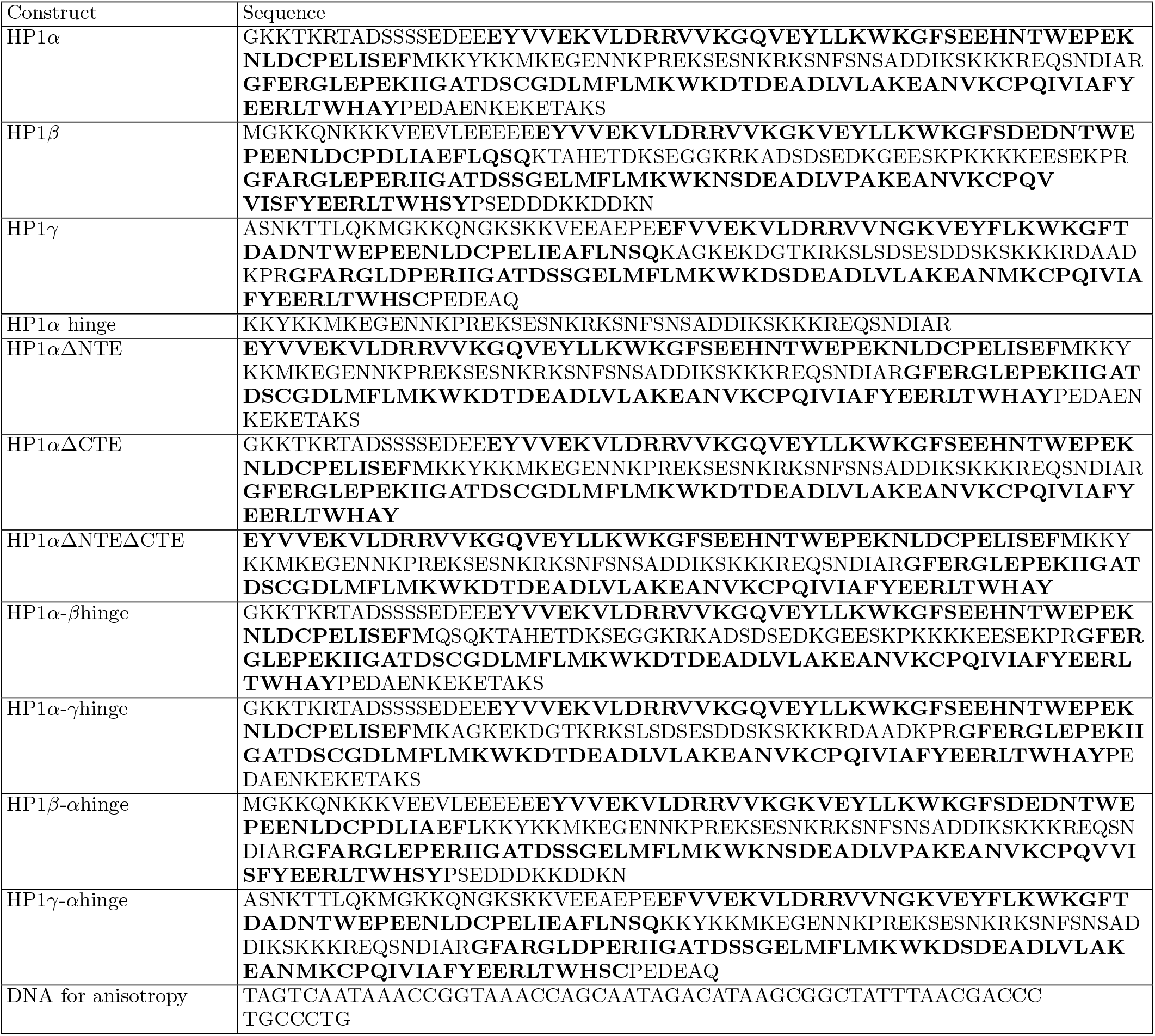
Protein sequences used in this study. Chromodomains (CD) and chromoshadow domains (CSD) are indicated in **bold**. A 6xHis tag followed by TEV cleavage site tag (MGHHHHHHDYDIPTTENLYFQGS) was appended to each construct for purification

**figure supplement 1.1.**
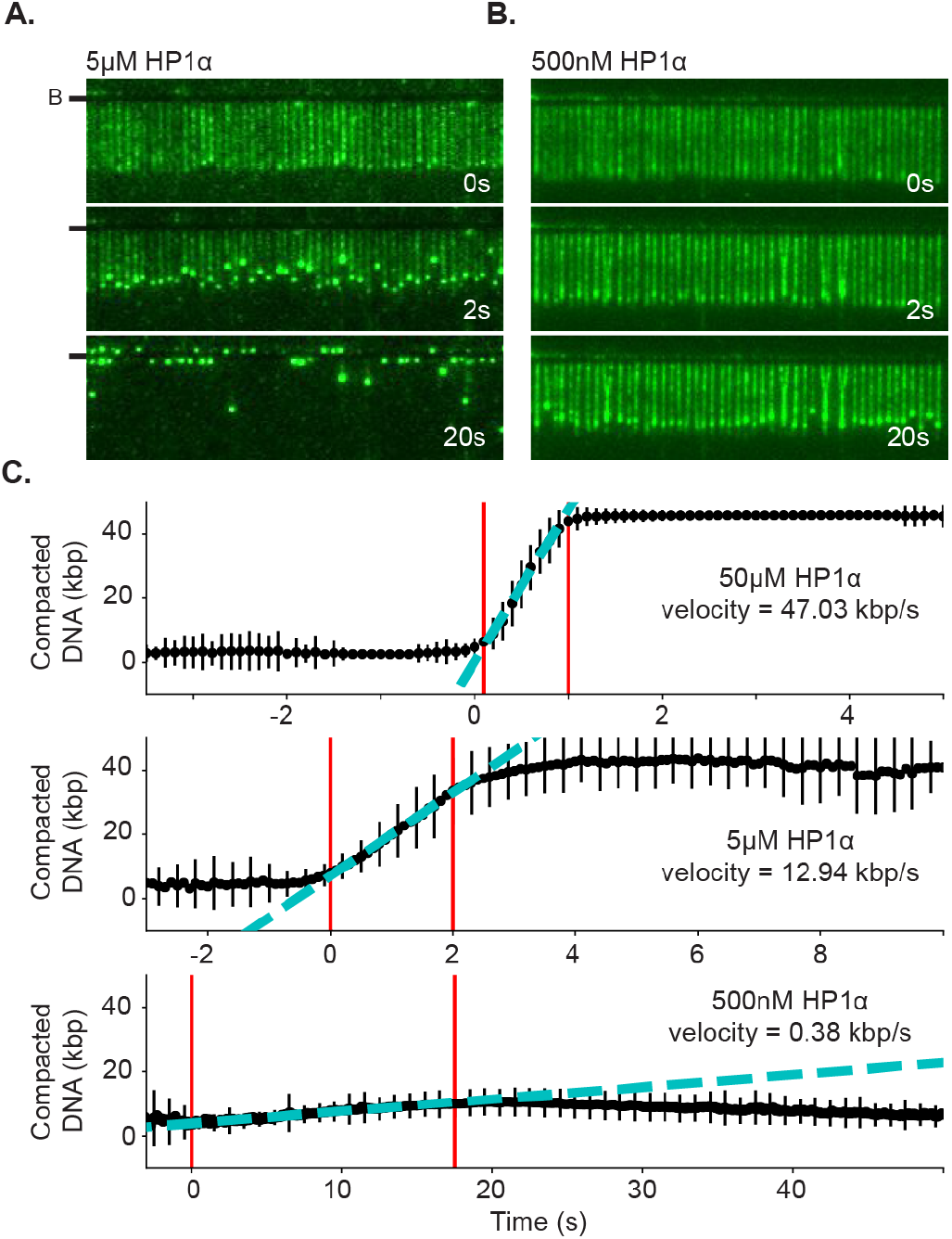
DNA compaction at different HP1*α* concentrations. **A.** and **B.** Timestamped images of DNA compaction by either **A.**5*µ*M or **B.**500nM HP1*α*. DNA is labeled with YOYO-1. (B-) or (-) specifies location of the barrier. **C.** Average DNA compaction for 50*µ*M, 5*µ*M and 500nM HP1*α*. Compaction velocity estimated from linear fit to data (cyan). Fit constrained to the region within the two red lines.

**figure supplement 1.2.**
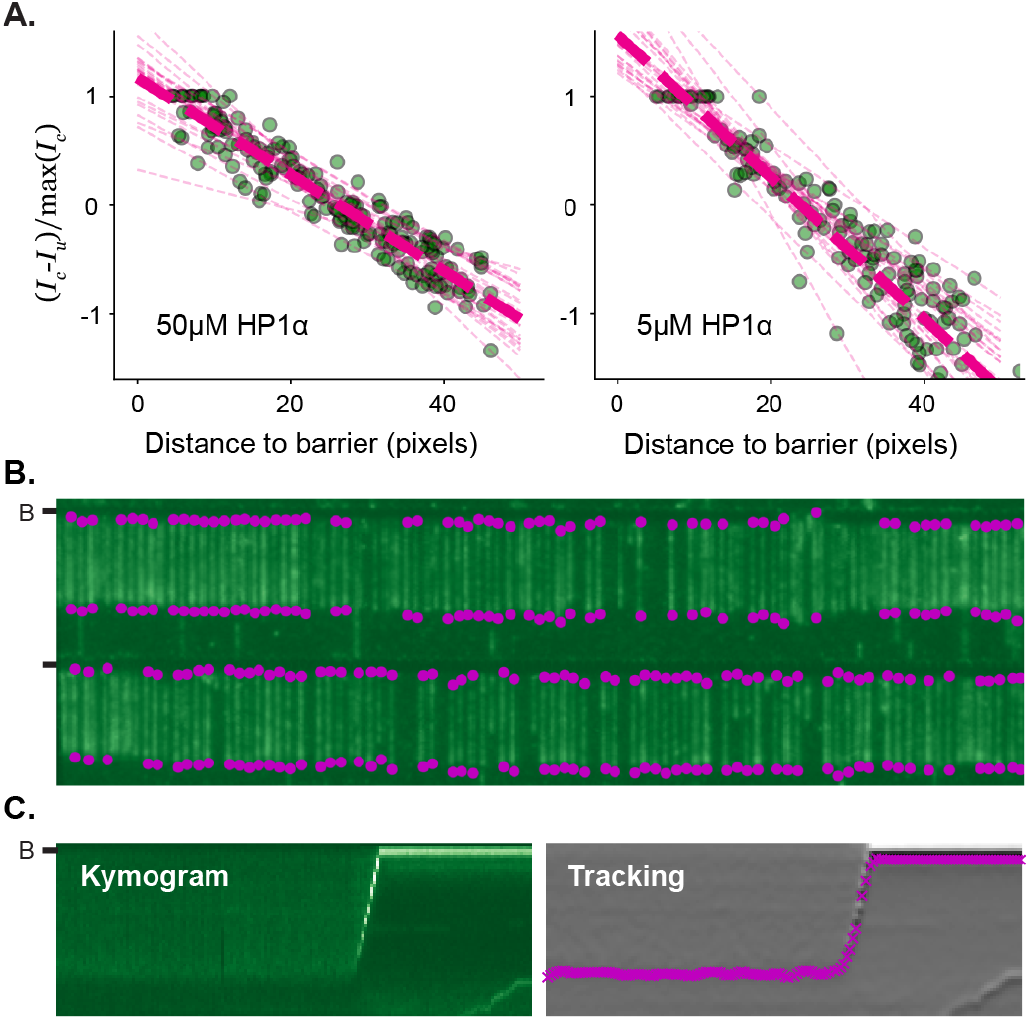
Fluorescence conservation and tracking DNA compaction by HP1*α*. **A.** Conservation of YOYO-1 fluorescence on DNA curtains. Normalized ratio of YOYO-1 intensity on the compacted and uncompacted segments. **B.** Automated detection of single DNA molecules within DNA curtains. **C.** Example output from tracking algorithm. (left) Kymogram showing compaction of DNA labeled with YOYO-1. (right) Overlay of tracking result on the derivative of the kymogram. (B-) or (-) specifies location of the barrier.

**figure supplement 2.1.**
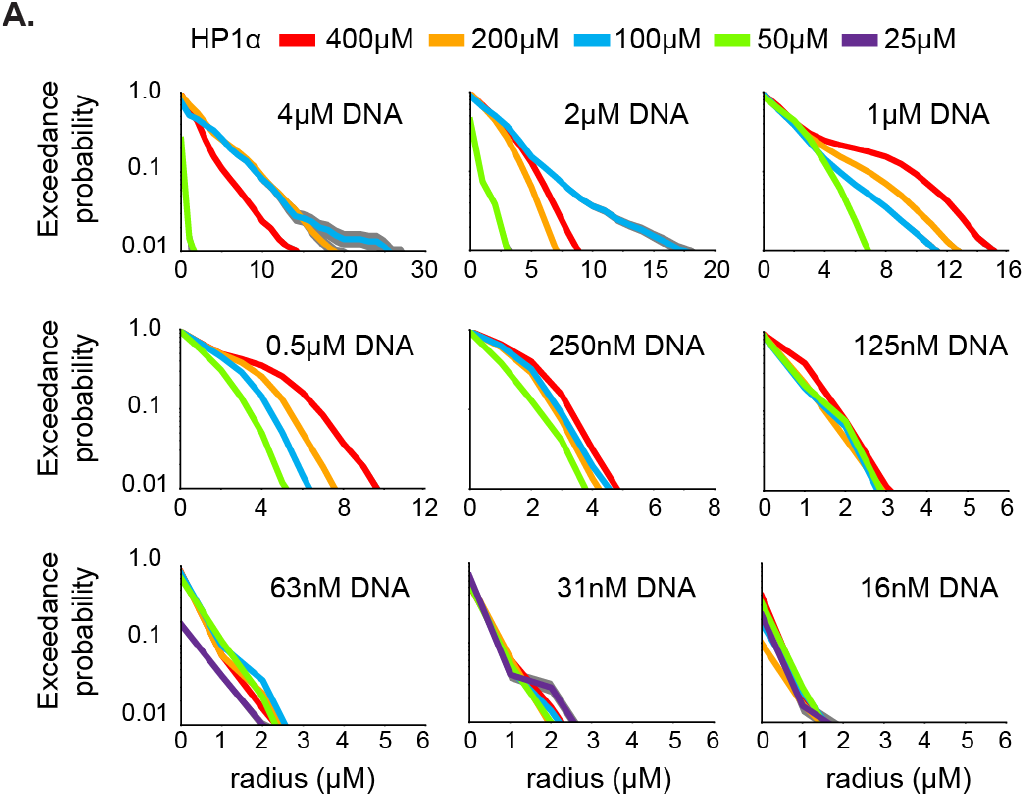
Exceedance probability. The number of condensates (y-axis) with radius exceeding indicated size (x-axis) for each concentration of HP1*α* and DNA in Figure 2A. Expectation values determined by integrating each curve are reported in Figure 2B-C.

**figure supplement 2.2.**
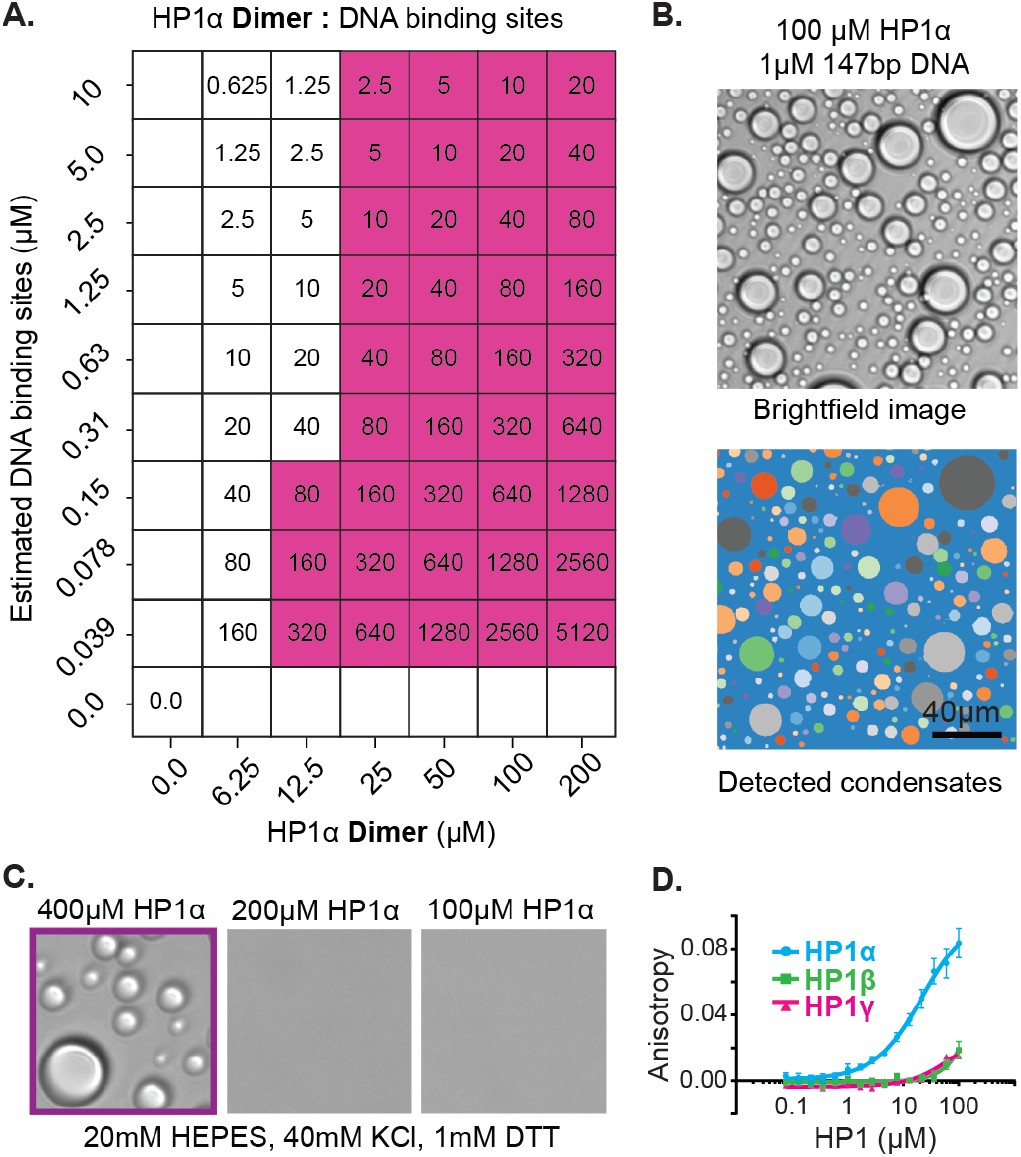
A. Characterization of HP1*α* condensates. **A.** Ratio of HP1*α* dimer to estimated DNA binding sites for experimental conditions in Figure 2A (2 HP1*α* binding sites per 147bp DNA oligo). **B.**(top) Brightfield image of 100*µ*M HP1*α* and 1*µ*M 147bp DNA and (bottom) output of automated condensate detection. **C.** Brightfield images of HP1*α* dialyzed into low salt buffer (20mM HEPES pH7.5, 40mM KCl, and 1mM DTT). **D.** Normalized fluorescence anisotropy curves for each HP1 paralog

**figure supplement 3.1.**
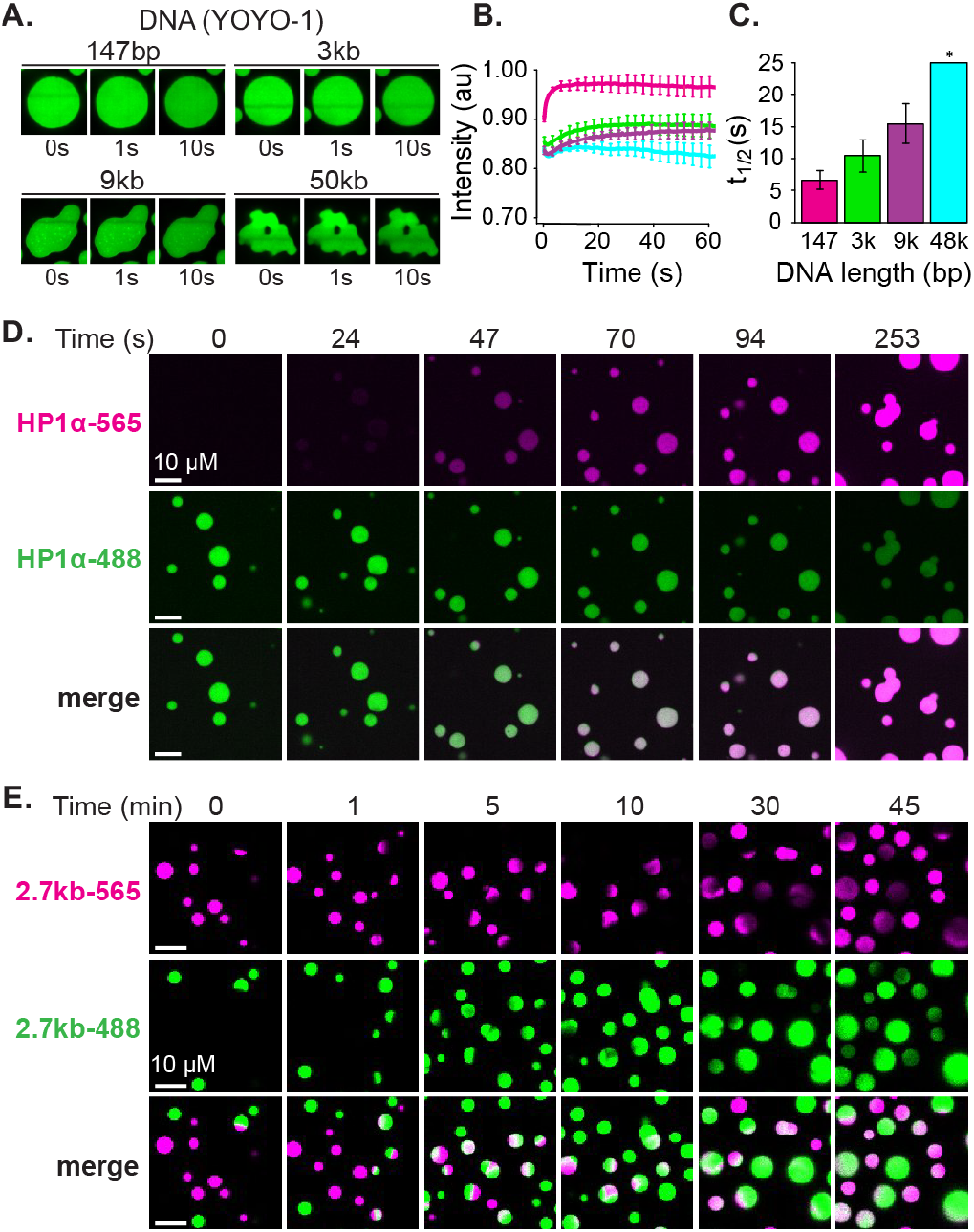
FRAP of DNA and mixing of HP1*α* and DNA in condensates. **A.** FRAP of YOYO-1 in condensates. Timestamped images from FRAP experiments for four lengths of linear DNA (147bp, 2.7kbp, 9kbp, or 50kbp). **B.** Recovery of YOYO-1 fluorescence intensity and **C.** half-life of recovery plotted for each DNA length tested. **D.** Timestamped images of two color HP1*α*-DNA condensate mixing experiments. Condensates formed separately with 2.7kbp unlabeled DNA and either HP1*α*-488 (green) or HP1*α*-565 (magenta). **E.** Timestamped images of two color HP1*α*-DNA condensate mixing experiments. Condensates formed separately with HP1*α* and 2.7kbp DNA-488 (green) or 2.7kbp DNA-565 (magenta).

**figure supplement 3.2.**
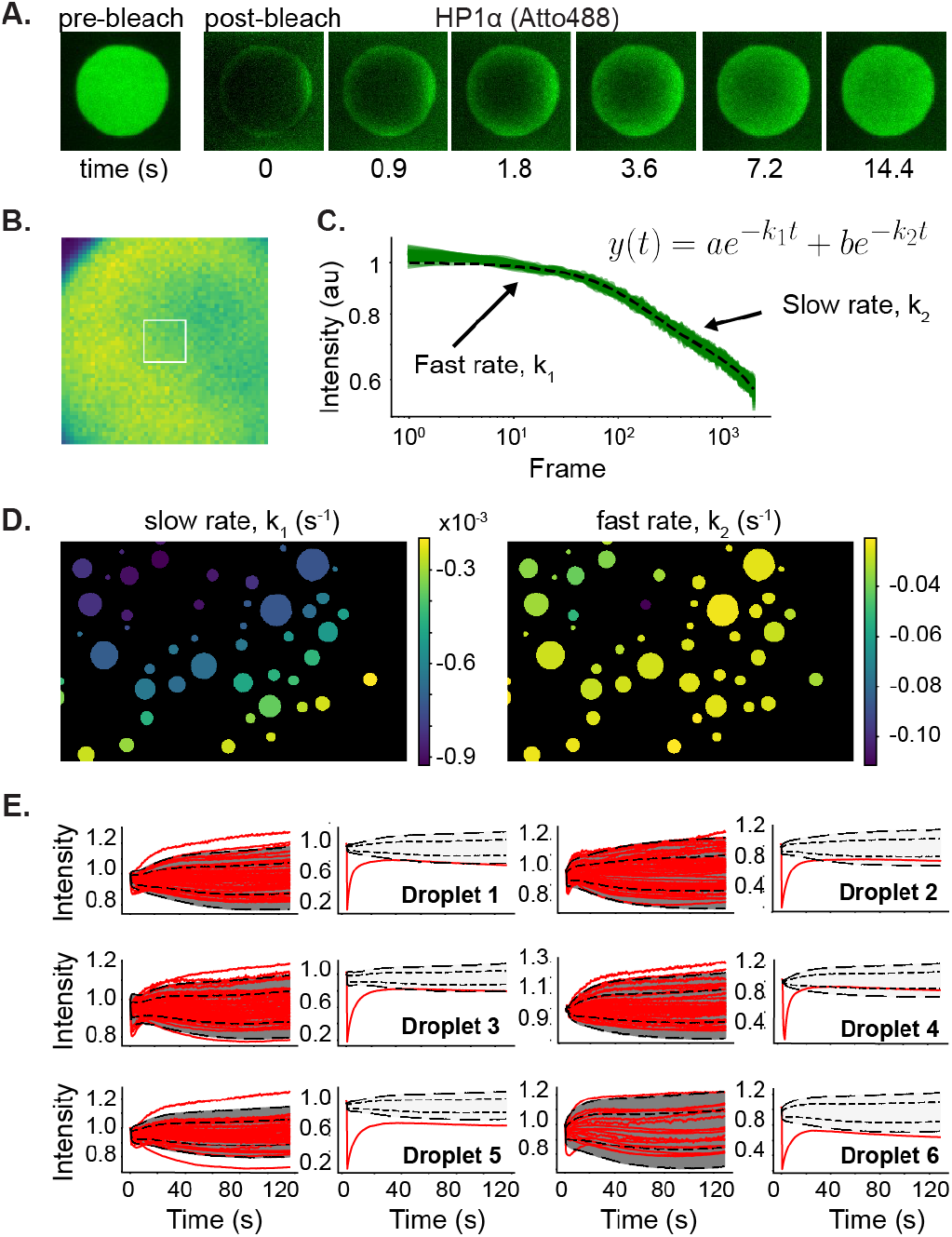
Whole droplet FRAP of HP1*α*-488 in HP1*α*-DNA condensates. **A.** Timestamped images of whole droplet HP1*α*-488 FRAP. **B.** and **C.** Time dependence of ambient HP1*α*-488 photobleaching within **B.** sample condensate region (white box), **C.** fit to a bi-exponential decay. **D.** Sample images colored by average fluorescence decay rates. **E.** Fluorescence recovery of ambient condensates (left) versus the photobleached condensate for six FRAP experiments. Dotted lines indicate 1 and 2 standard deviations from the mean determined from the ambient condensates.

**figure supplement 4.**
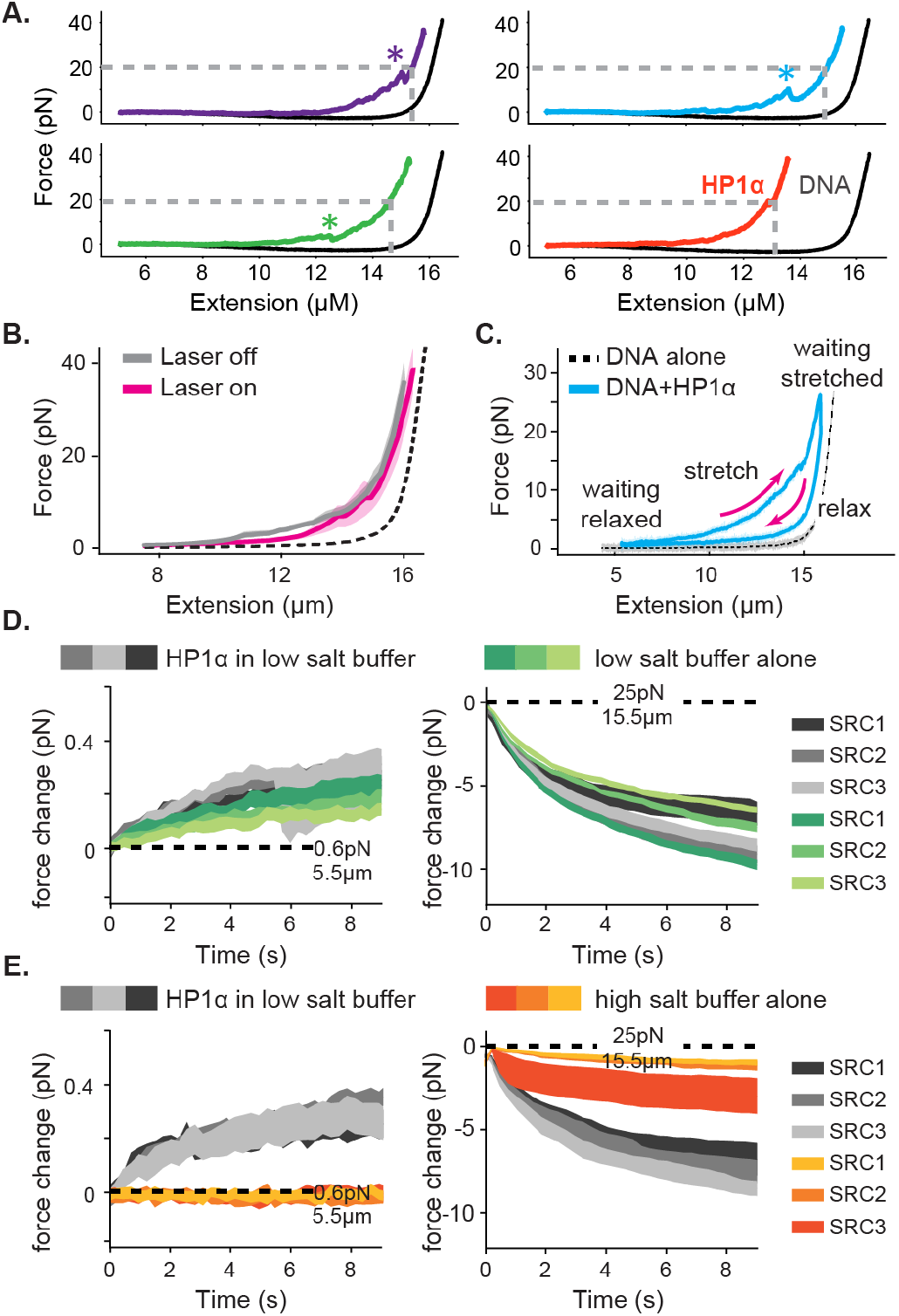
Representative traces and controls for optical trap experiments. **A.** Four representative traces from Figure 4C. All traces are separate pulls from the same DNA strand. **\*** indicates rupture event. Grey dashed line indicates the DNA extension at 20pN force reported in Figure 4D. **B.** Average force extension curves for the second SRC either with (magenta) or without (gray) laser illumination. The force extension curve of DNA alone is shown in black. **C.** Force extension curve across a stretch-relax cycle including waiting periods in the extended or relaxed configurations. **D.** and **E.** Force change in the relaxed (left) and stretched (right) configurations in the presence (gray) and absence (green) of HP1*α*. SRCs in the absence of protein performed in either **D.** low salt (70mM KCl) or **E.** high salt (500mM KCl) buffer.

**figure supplement 5.**
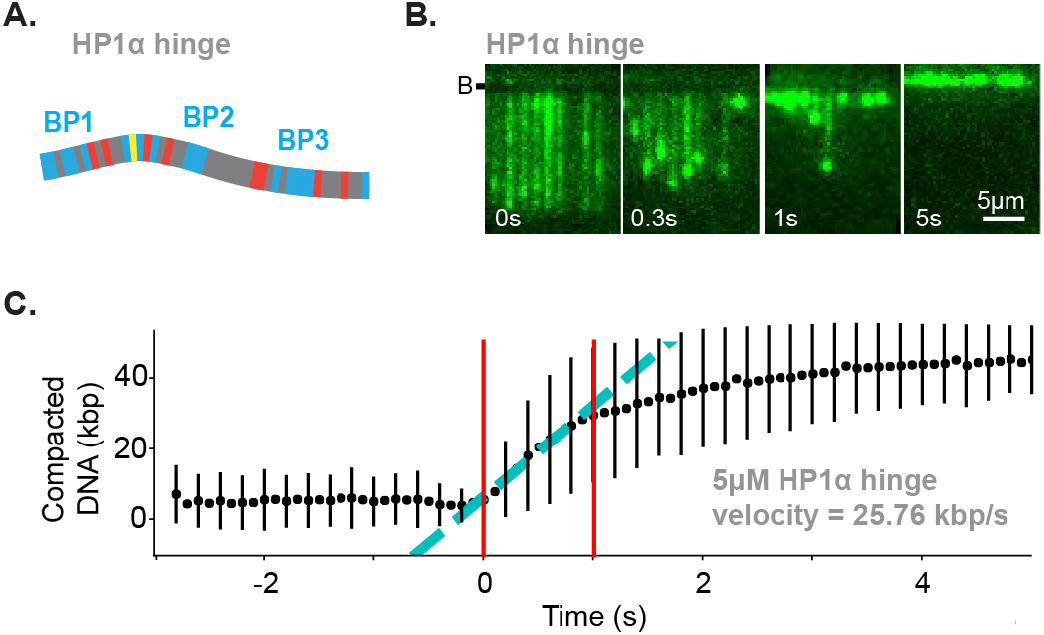
The hinge region of HP1 is sufficient for DNA compaction. **A.** Cartoon of HP1*α* hinge with color-coded disordered residues: positive residues (K and R) blue, negative residues (E and D) red, proline yellow, and all other residues grey. The HP1*α* hinge contains three basic patches (BP). **B.** Timestamped images of DNA labeled with YOYO-1 undergoing compaction by 5*µ*M HP1*α* hinge (unlabeled) shown before, during, and after compaction. (B-) specifies location of the barrier. **C.** Average DNA compaction by the HP1*α* hinge. Compaction velocity estimated from linear fit to data (cyan). Fit constrained to the region within the two red lines.

**figure supplement 6.1.**
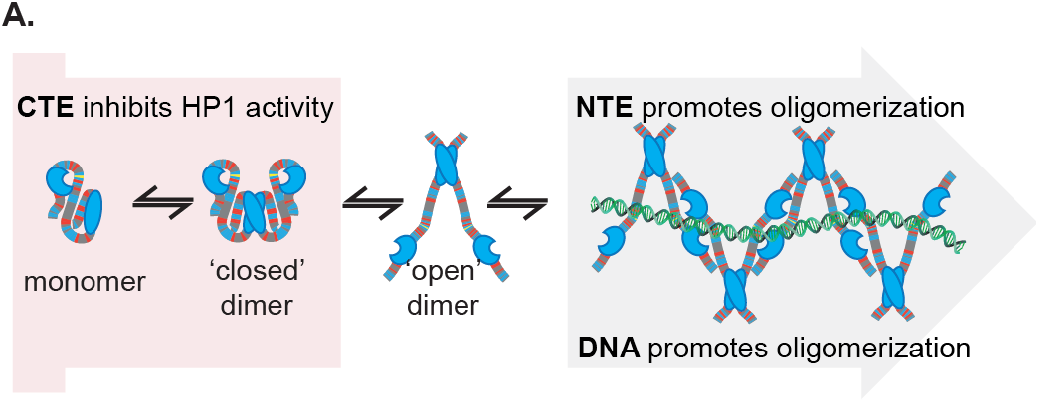
A. Proposed model of HP1*α* autoregulation and potential oligomerization.

**igure supplement 6.2.**
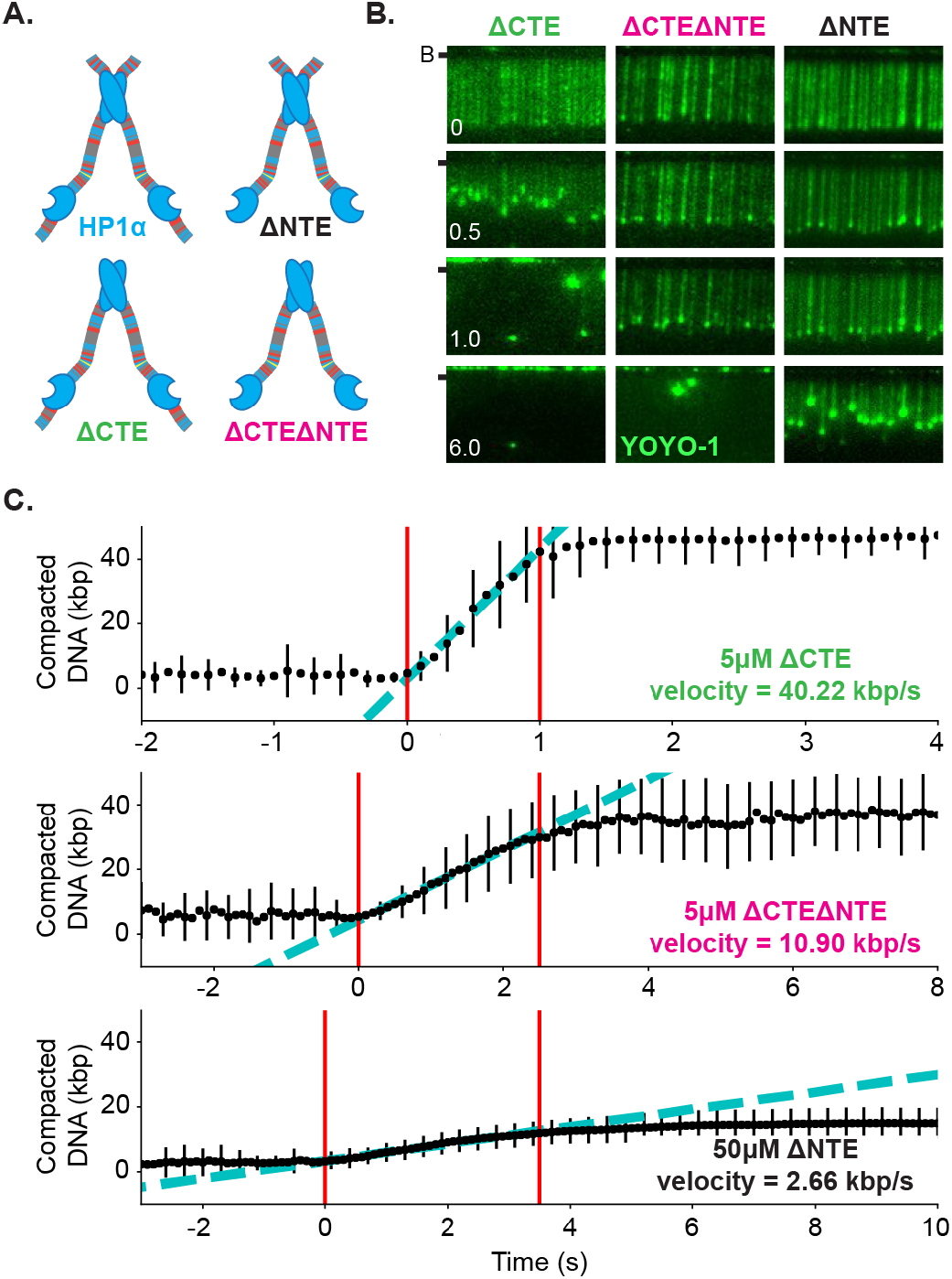
DNA compaction activity of HP1*α* domain mutants. **A.** Cartoon of HP1*α* extension mutants with color-coded disordered residues: positive residues (K and R) blue, negative residues (E and D) red, proline yellow, and all other residues grey. **B.** Timestamped images of DNA labeled with YOYO-1 undergoing compaction by 5*µ*M HP1*α*ΔCTE, 5*µ*M HP1*α*ΔNTEΔCTE, and 50*µ*M HP1*α*ΔNTE (unlabeled) shown before, during, and after compaction. (B-) or (-) specifies location of the barrier. **C.** Average DNA compaction by each HP1*α* mutant. Compaction velocity estimated from linear fit to data (cyan). Fit constrained to the region within the two red lines.

**figure supplement 7.**
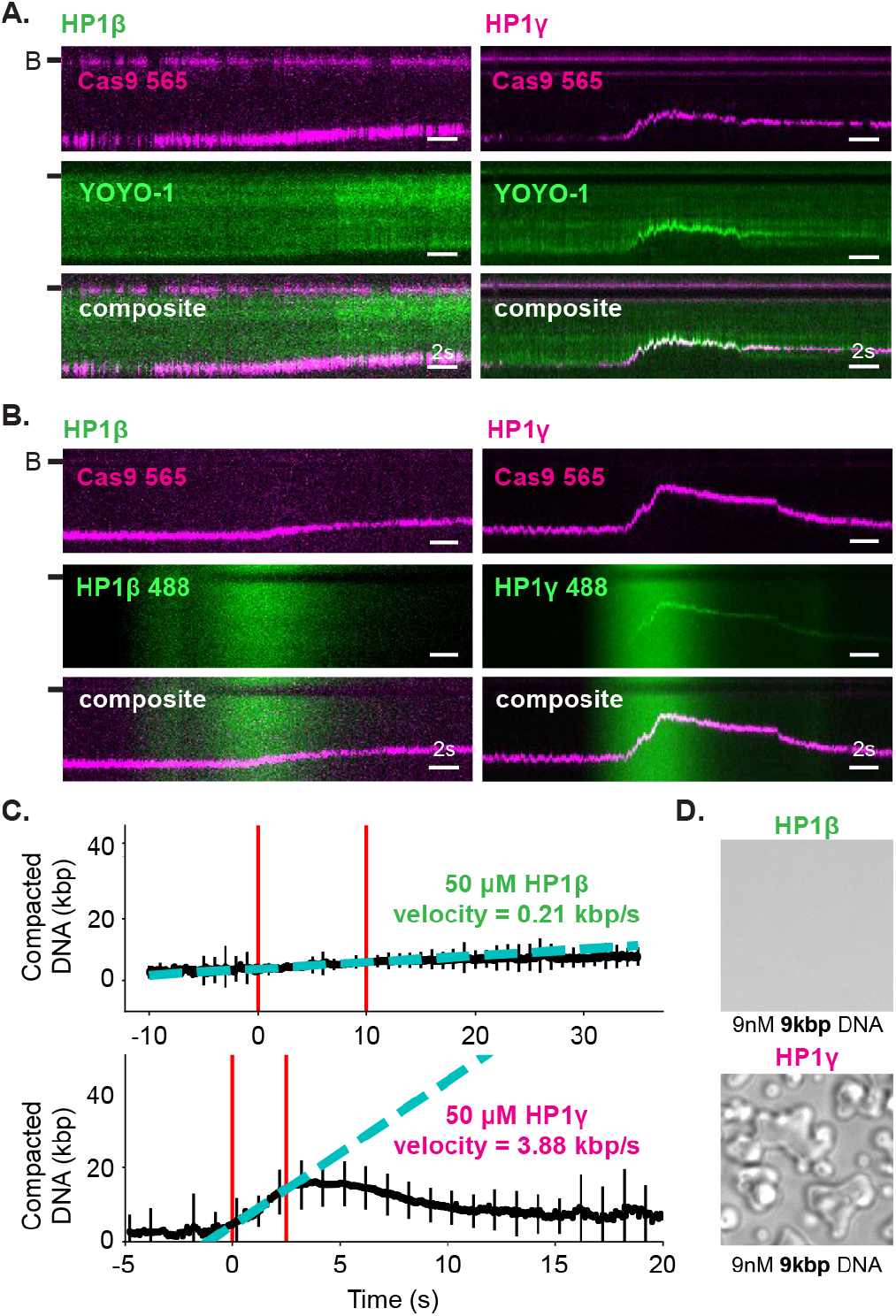
DNA compaction by HP1*β* and HP1*γ*. **A.** and **B.** Kymograms of DNA compaction by 50*µ*M HP1*β* (left) and HP1*γ* (right). **A.** DNA labeled with YOYO-1 (top), dCas9-565 (middle), and composite image (bottom). **B.** HP1*α*-488 (top), DNA labeled with dCas9-565 (middle), and composite image (bottom). (B-) or (-) specifies location of the barrier. **C.** Average DNA compaction by HP1*β* (top) and HP1*γ* (bottom). Compaction velocity estimated from linear fit to data (cyan). Fit constrained to the region within the two red lines. **D.** Brightfield images of HP1*β* (top) and HP1*γ* (bottom) and 9kbp DNA.

## Notes

### Competing Interest Statement

The authors have declared no competing interest.

